# Senescent cell networks link matrix remodeling and vascular dysfunction in human fibroids

**DOI:** 10.64898/2026.08.06.743362

**Authors:** Joscelyn C. Mejías, Nazmiye Celik, Sushma Nagaraj, Katlin B. Stivers, Helen H. Nguyen, A. Shri Ramanujam, Frank Haoning Yu, Maria A. Browne, Rachel Michel, Md Soriful Islam, Christopher Cherry, Alexandra N. Rindone, Abigail Fennell, Chanhong Min, Bhuchitra Singh, Kavita Krishnan, Anna Ruta, Natalie Rutkowski, Malak El Sabeh, Sadia Afrin, Yiting Chen, Samya El Sayed, Pei-Hsun Wu, Jude M. Phillip, Elana J. Fertig, Mostafa A. Borahay, James Segars, Jennifer H. Elisseeff

## Abstract

Uterine fibroids (leiomyomas) are highly prevalent benign tumors defined by excessive extracellular matrix (ECM) deposition, altered vascular structure, and progressive tissue stiffening, yet the cellular programs that coordinate these features remain poorly understood. Cellular senescence has been implicated in fibroid biology, but whether senescence represents a uniform state or distinct, functionally specialized cell identities within fibroids is unknown. Here, we identify the distinct heterogeneous populations of senescent cells (“senotypes”) present in human fibroids and characterize their role in shaping the fibroid microenvironment. Using single-cell RNA sequencing (scRNA-seq) integrated with a senescence gene signature and protein-level validation, we identify senescent cells (SnC) distributed across fibroblast, mural, and endothelial compartments, each exhibiting distinct transcriptional programs. SnC endothelial cells (ECs) are enriched in fibroids relative to matched myometrium and activate *TEAD4*-associated mechanosensing, angiogenic, and immune signaling pathways, despite being associated with impaired vessel maturation *in situ*. In parallel, SnC fibroblast and mural populations in fibroids upregulated SRF-associated cytoskeletal and ECM programs, accompanied by increased *COL6A3* expression and collagen VI deposition, consistent with tissue stiffening. Ligand–receptor and spatial analyses reveal that these SnC populations function as interconnected signaling hubs, coordinating immune cell recruitment and stromal remodeling. Importantly, analysis of human fibroids treated with collagenase demonstrated a reduction in both ECM density and SnC burden, supporting a reinforcing relationship between matrix mechanics and senescence. Together, these findings establish senescence in fibroids as a heterogeneous, mechanically reinforced, and network-driven process that links vascular dysfunction, immune signaling, and fibrosis, highlighting distinct SnC states as potential translational targets for non-surgical therapies.

## Introduction

Uterine fibroids (leiomyomas) are among the most common tumors in women, affecting up to 70–80% of individuals over their lifetime and representing a leading cause of hysterectomy worldwide [1–3]. Though benign, fibroids impose a substantial clinical burden on millions of women, causing heavy menstrual bleeding, anemia, pelvic pain, infertility, and pregnancy complications, and generating significant healthcare costs [3–7]. A defining feature of fibroids is their dense, collagen-rich ECM coupled with altered vascular architecture and progressive tissue stiffening, hallmarks that align fibroids with fibrotic diseases more broadly [8–10]. However, unlike many fibrotic pathologies [11–13], the cellular and microenvironmental programs that sustain this remodeling remain incompletely defined.

Recent transcriptomic and histological studies have revealed that fibroids are not homogeneous masses of smooth muscle cells, but instead comprise diverse stromal, vascular, and immune populations that contribute to matrix deposition, altered vascular function, and tissue organization [14, 15]. These observations suggest that fibroid progression emerges from coordinated interactions across cell types rather than a single dominant lineage. However, a unifying framework that explains how these cellular compartments integrate to drive fibrosis, vascular dysfunction, and tissue mechanics is lacking [16].

Cellular senescence has emerged as a central regulator of tissue remodeling across aging, injury, and fibrosis [17, 18]. Classically defined as a stable cell-cycle arrest marked by expression of p16INK4a, p21(CDKN1A), and senescence-associated β-galactosidase, SnCs are now recognized as functionally active and highly context-dependent states [19]. Through the senescence-associated secretory phenotype (SASP), these cells secrete cytokines, growth factors, and matrix-remodeling enzymes that can reshape tissue architecture and intercellular communication [20]. Importantly, senescence is not a singular identity but rather encompasses diverse phenotypic states that depend on cell type, microenvironment, and mechanical context [20–22].

SnCs have been reported to accumulate in fibroids [23, 24], as well as in aged [25] and injured myometrium [26, 27]. However, what defines a senescent cell in fibroids, and whether distinct SnC types perform specialized functions, remains unclear.

Emerging evidence across fibrotic diseases indicates that SnCs can adopt lineage-specific roles, including promoting fibrosis, regulating vascular remodeling, and directing immune responses [22, 28, 29]. Thus, understanding senescence in fibroids requires moving beyond marker-based identification toward defining SnC identity (“senotypes”) and their functional roles within tissue networks. A second unresolved question is how senescence interfaces with the defining features of fibroids, including matrix accumulation, vascular dysfunction, and immune engagement. Fibroids exhibit paradoxical vascular phenotypes, including reduced vessel density despite persistent angiogenic signaling [10, 30], together with increased tissue stiffness [8]. In parallel, immune cells are present within fibroids but are spatially and functionally distinct from normal myometrium [14, 15, 31, 32]. These features suggest that fibroids arise from coupled vascular–stromal–immune interactions, potentially coordinated by SnC populations. However, whether senescence acts as a central organizing principle linking these processes has not been established.

Here, we address these gaps by defining the identity, distribution, and functional roles of SnC in human fibroids. Using single-cell RNA sequencing (scRNA-seq) integrated with a senescence gene signature and transfer learning approaches, we map senescent cell states across stromal, vascular, and immune compartments in fibroids and matched myometrium. We identify multiple senotypes with distinct transcriptional and signaling programs, including endothelial, fibroblast, and mural populations linked to vascular remodeling, ECM production, and immune communication. We further show that these senescent cell states engage in networked signaling interactions that contribute to remodeling the fibroid microenvironment, and that these programs are tightly coupled to mechanosensitive transcriptional pathways and tissue stiffness. Finally, leveraging human clinical samples from a collagenase treatment study, we demonstrate that matrix remodeling reduces SnC burden, providing translational evidence linking ECM mechanics and senescence in fibroids.

Together, this work defines a framework in which heterogeneous SnC states coordinate vascular dysfunction, immune engagement, and matrix remodeling within fibroids. This framework positions senescence as both a mechanistic driver of fibroid pathogenesis and a potential therapeutic target for non-surgical intervention.

## Results

### Senescent cells are enriched and heterogeneous in uterine fibroids

Uterine fibroids and patient-matched myometrium samples were collected from patients undergoing either elective hysterectomy or myomectomy (30 patient samples were collected for multiple analyses across the experiments, as listed in **Supplementary Table 1**). To determine whether SnCs are enriched in fibroids, expression of the senescence-associated marker p16 [33] was assessed at the protein level in fibroid and matched myometrium samples (**Fig. 1a,b**).

**Figure 1.**
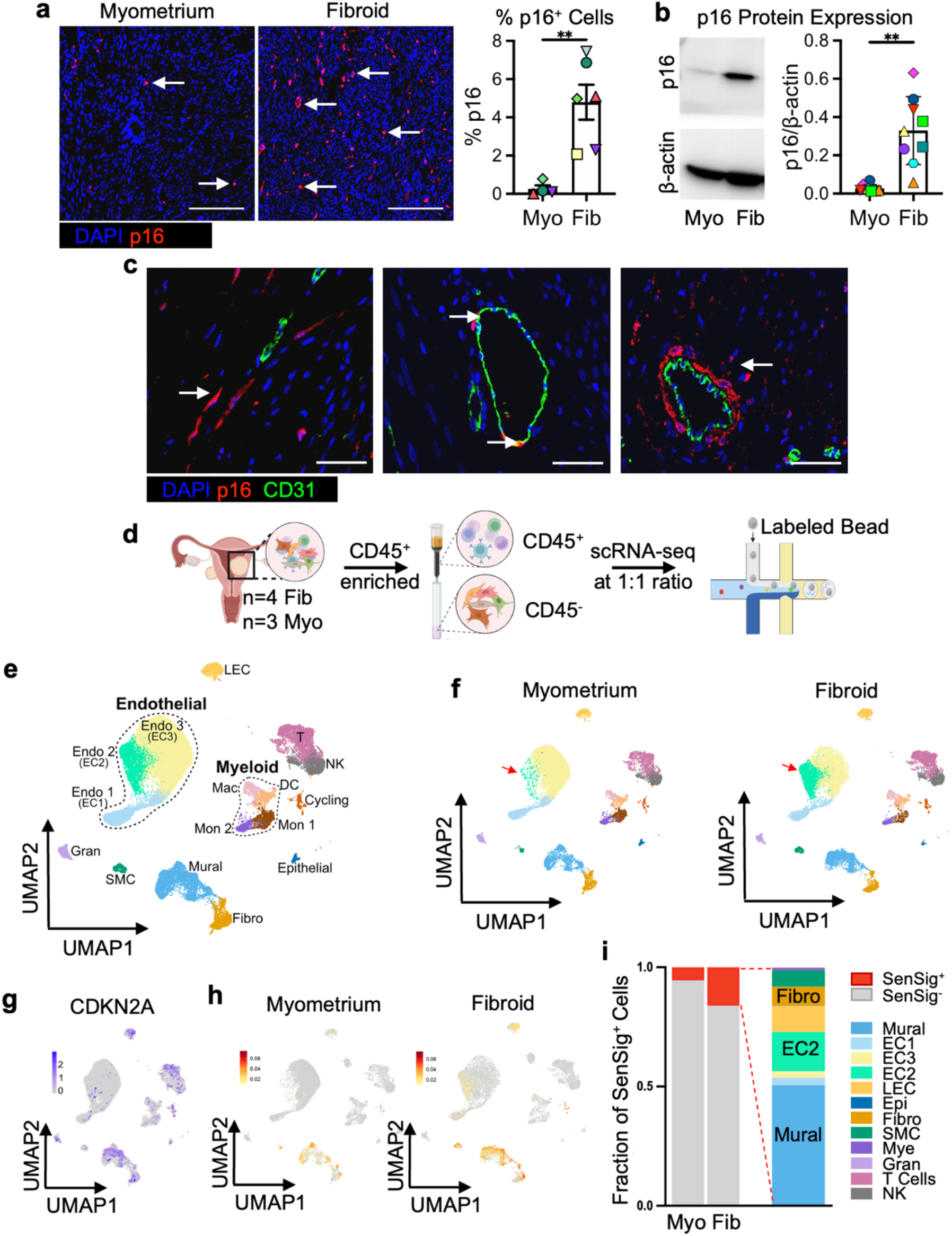
Uterine fibroids contain expanded and diverse SnC niches. **(a)** IF imaging and quantification of DAPI (nuclei, blue) and p16 (senescence, red) in a FFPE section of myometrium (Myo) and fibroid (Fib). Scale bar, 200 μm; arrows indicate cells of interest (*n* = 6 independent samples). **(b)** Representative western blot (WB) and quantification. Each patient is given a unique color, shape combination in the quantifications (*n* = 9 paired samples). **(c)** p16 SnC cell heterogeneity observed across IF of DAPI (nuclei, blue), p16 (senescence, red), and CD31 (vessels, green) in fibroid samples; scale bar = 50μm, white arrows identify cells of interest. **(d)** Schematic overview of the scRNA-seq workflow. **(e)** Visualization of uniform manifold approximation and projection (UMAP) showing 16 high-quality cells clusters from human myometrium (*n* = 3), fibroid (*n* = 4) tissue together. **(f)** UMAP projections split by tissue type, highlighting differences in cellular composition between fibroid and myometrium samples. Red arrows indicate one endothelial cell cluster (EC2), which is enriched in fibroid tissues. **(g)** Feature plot showing expression of *CDKN2A* (p16) across the scRNA-seq dataset. **(h)** UMAP projection overlaid with SenSig transfer-learning scores, with SenSig^+^ cells highlighted and SenSig^-^ cells shown in grey, displayed separately for fibroid and myometrium. **(i)** Bar plots showing the relative abundance of SenSig^+^ cells in myometrium and fibroid tissues (left) and the contribution of individual cell types to the senescent cell pool within fibroids (right). a-b normality tested with Shapiro-Wilk test followed by Welch’s t test for **(a)** and Wilcoxon test for **(b)**, significance determined using p<0.05 for each test. Statistical significance: **, *P* < 0.01.

Immunofluorescence (IF) analysis demonstrated a significant increase in p16⁺ cells in fibroid sections relative to matched myometrium (**Fig. 1a**). Quantification across tissue sections confirmed this enrichment (4.8 ± 2.24% versus 0.27 ± 0.35%, respectively; **Fig. 1a**). Consistent with these findings, western blot (WB) analysis revealed an approximately 11-fold increase in p16 protein levels in fibroids (**Fig. 1b**, **Supplementary Fig. 1a**).

In addition to increased abundance, p16 IF staining revealed substantial morphological and spatial heterogeneity of SnCs within fibroid tissue (**Fig. 1c**). In stromal regions, p16⁺ cells frequently exhibited elongated morphologies consistent with fibroblast-like identities. p16 expression was also observed adjacent to CD31⁺ vascular structures, consistent with mural cell localization, and co-localization of p16 with CD31 indicated the presence of SnC ECs (**Fig. 1c** and **Supplementary Fig. 1b**). These findings indicate that SnCs span multiple cellular compartments and are spatially organized within distinct niches in fibroids.

To characterize this heterogeneity at transcriptional resolution, scRNA-seq was performed on fibroid (n = 4) and matched myometrium (n = 3) samples (**Fig. 1d**). Given the dense ECM of fibroids, tissue dissociation conditions were varied and found to influence the cell types recovered. Therefore, we tested four different tissue dissociation conditions and selected an optimized digestion protocol that ensured recovery of diverse stromal (CD45^-^) and immune (CD45^+^) populations prior to scRNA-seq (**Supplementary Fig. 2**). To improve immune-cell representation, CD45^+^ cells were enriched to achieve a 1:1 CD45^+^: CD45^-^ cell ratio. Following quality control, removal of low-quality cells, and batch correction (**Supplementary Fig. 3a,b**), a final dataset of 41,159 cells was obtained and cluster into 15 transcriptionally distinct stromal and immune populations (**Fig. 1e**, **Supplementary Fig. 4** and **Supplementary Table 2**). Stromal populations included ECs (EC1, EC2 and EC3) *(VWF*, *PECAM1)* (**Supplementary Figs. 4, 5**), smooth muscle cells (SMC) (*MYH11*, *ACTA2*), mural cells (*RGS5*, *ACTA2)*, fibroblasts (*COL1A1*, *DCN*), and lymphatic endothelial cells (LEC) (*LYVE1*, *PDPN*), while immune populations comprised T cells (*CD3E*), natural killer (NK) cells (*NKG7*), granulocytes (Gran) (*KIT*), and myeloid cells (*ITGAX*, *CD68*) (Supplementary Fig. 4). Myeloid subclusters include monocytes (Mon1 and Mon2), dendritic cells (DC), and macrophages (Mac) (**Supplementary Figs. 4,6**). A cycling cluster (*MKI67*, *TOP2A*) was also identified (**Supplementary Fig. 4)**.

Comparison of tissue-specific cell type abundances using edgeR while controlling for patient variability revealed shifts in cellular composition between fibroids and myometrium, including enrichment of the EC2 endothelial population in fibroids (edgeR’s glmQLFTest [34], FDR = 0.0004) (**Fig. 1f**, **Supplementary Fig. 7a-b** and **Supplementary Table 3**). While these analyses defined the cellular landscape, further analysis of the scRNA-seq data is needed to resolve SnCs across cell types.

### SenSig**–**projectR identifies senescent cell states across stromal compartments

To identify senotypes within the scRNA-seq dataset, the expression of canonical SnC-associated marker genes was first evaluated. *CDKN2A (p16)* expression was detected in subsets of stromal populations, including EC2, mural, and fibroblast clusters (**Fig. 1g**), consistent with the detection of p16 in these the protein-level IF data (**Fig. 1c**). However, *CDKN2A* expression is not exclusive to SnCs and can also be influenced by cellular stress and cell-state heterogeneity [35, 36]. In addition, transcript dropout inherent to scRNA-seq can limit reliable detection of SnC-associated genes [37]. Other genes, such as *CDKN2B* (*p15*), *CDKN1A* (*p21*), and SASP genes such as *IL6*, were expressed in a non-specific manner across all the clusters (**Supplementary Fig. 8a**). Together, these findings indicate that individual markers are insufficient to identify SnCs at transcriptional resolution.

To overcome the limitations posed by defining SnCs from expression values of individual marker genes, we applied our previously established *in vivo* senescence gene signature (SenSig) [38] using the transfer learning tool projectR [39, 40]. SenSig is a SnC-associated gene signature derived from *in vivo* senescent cell states. Using SenSig, we performed a generalized least squares fit to the scRNA-seq dataset implemented in the projectR package. Cells with projection weights significantly greater than zero (*p* < 0.01) were classified as senescent (SenSig^+^) (**Fig. 1h**, **Supplementary Fig. 8b** and **Supplementary Table 4**). SMC, mural, fibroblast, LEC, EC2, and cycling clusters exhibited higher projection weights (**Supplementary Fig. 8b**). We excluded the cycling cluster because of its high expression of proliferative markers, including *TOP2A* and *MKI67* (**Supplementary Fig. 4b**). We sought to define the genes distinguishing the SnC programs across the stromal populations using a statistic that prioritizes genes that distinguish the drivers of projection weights between sample groups called projectionDrivers [41]. SnC fibroblasts were enriched for ECM-remodeling genes including *TNC*, *COL11A1*, and *FBLN7*, whereas SnC mural cells showed ECM-remodeling and inflammation-associated genes including *COL23A1*, *MMP11*, *IL34*, and *SERPINE2*. SnC EC2 cells were associated with SASP- and endothelial activation-associated genes including *ANGPTL4*, *SPP1*, *UNC5B*, and *CD276*. *FJX1* was shared across multiple stromal populations (**Supplementary Fig. 8c**).

SenSig⁺ cells demonstrated an increase in transcriptionally defined SnCs in fibroids relative to myometrium, comprising approximately 16% of fibroid cells compared to approximately 5% in myometrium (**Fig. 1i** and **Supplementary Table 5**). SnCs were identified predominantly in mural, fibroblast, and EC2 populations (**Fig. 1i**). SnC LECs and SMCs were excluded from further analysis due to limited representation (LECs were detected in only one patient, and SMCs had low cell counts) (**Supplementary Fig. 9**). Together, these findings demonstrate that transcriptionally defined SnCs are expanded in fibroids and concentrated within defined stromal compartments, including mural, fibroblast, and EC2 populations (**Fig. 1i**), consistent with protein-level validation of p16 (**Fig. 1c**). We next examined the functional characteristics of these senescent states.

### Distinct senescent cell states exhibit specialized SASP and receptor programs

To define functional differences between SnC and non-SnC populations, we performed differential gene expression analysis between SenSig⁺ (SnC) and SenSig⁻ (non-SnC) cells within the fibroblast, mural, and EC2 populations, each containing more than 200 cells per group (**Fig. 2a**, **Supplementary Fig. 10** and **Supplementary Tables 6–8**). Because SnCs exert their effects through the SASP while also responding to microenvironmental cues, we examined differentially expressed genes encoding both secreted ligands and cell-surface receptors within SenSig⁺ populations (adjusted *P* value < 0.05, avg_log2FC > 0.3). To characterize these features, differentially expressed genes (DEGs) were annotated using MatrisomeDB [42], Surfaceome [43], and CellPhoneDB [44] to identify matrix-associated factors, secreted factors (SASP), and cell-surface proteins (**Fig. 2a**, **Supplementary Fig. 10** and **Supplementary Table 9**).

**Figure 2.**
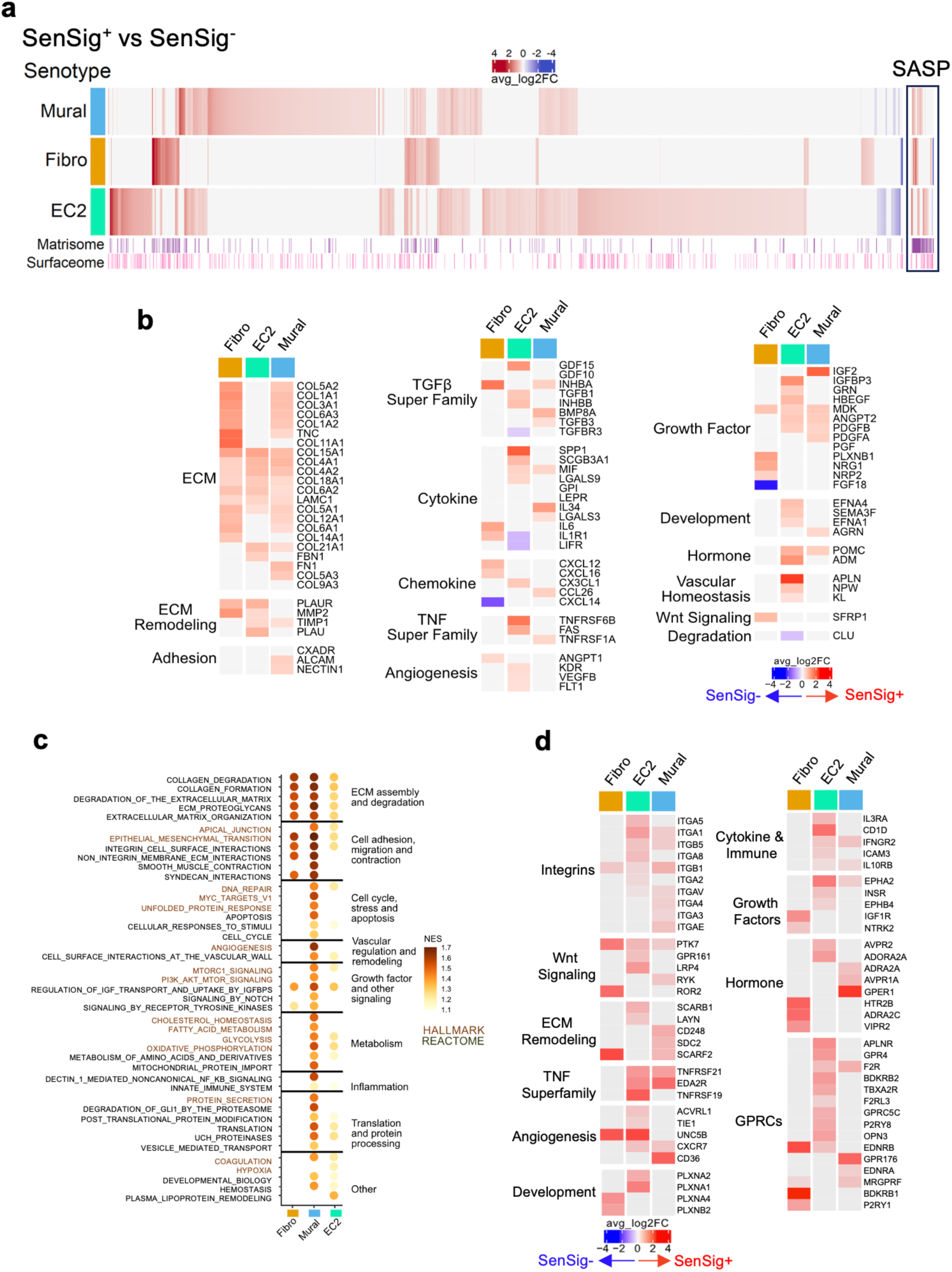
**Unique SASP and receptor signatures characterize vascular and fibrotic SnC phenotypes**. **(a)** Heatmap showing DEG between SenSig⁺ and SenSig⁻ cells within mural, fibroblast, and EC2 clusters, with annotation of matrisome- and surfaceome-associated genes (SASPs represent secreted factors). Colors indicate the average log₂ fold change (avg_log₂FC), with red denoting higher expression in SenSig⁺ cells and blue denoting higher expression in SenSig⁻ cells. **(b)** Heatmaps of selected DEG encoding secreted factors associated with cell-type-specific SASP programs across Fibro, EC2, and mural populations. Genes are organized according to their principal functional categories, including extracellular matrix organization and remodeling, cytokine and chemokine signaling, growth-factor signaling, angiogenesis, development, and vascular homeostasis. **(c)** Summary of selected pathways from gene set enrichment analysis (GSEA) of SenSig⁺ vs SenSig⁻ cells revealing functional pathways. **(d)** Heatmaps of selected DEG encoding non-secreted cell-surface receptors across Fibro, EC2, and mural populations. Receptors are grouped by functional category, including integrins, cytokine and immune receptors, growth-factor receptors, hormone receptors, and G protein–coupled receptors (GPCRs). Color direction is as described in **(a)**.

Analysis of SASP factors showed enrichment of secreted ligands associated with ECM organization and remodeling across SnC populations (**Fig. 2b**). We identified both shared and distinct SASP programs across SnC types (**Supplementary Fig. 11a**). Shared upregulated SASP factors included multiple collagens (*COL6A1, COL6A2, COL4A2, COL4A1, COL15A1, COL18A1, COL5A1,* and *COL21A1*), additional ECM-associated proteins (*LAMC1* and *TNC*), and the growth factor *MDK*. These matrix-associated SASP factors were more prominently increased in SnC mural and fibroblast populations relative to their non-SnC counterparts (**Fig. 2b** and **Supplementary Fig. 11a**). In addition to these shared factors, SnC populations expressed SASP components from the TGF-β and TNF superfamilies, as well as cytokines and chemokines including *IL6*, together with angiogenic and growth-associated factors (**Fig. 2b**). Although most secreted ligands were upregulated in SenSig⁺ cells, *CXCL14* and *FGF18* were downregulated in SnC fibroblasts, whereas *TGFBR3, LIFR, CLU*, and *IL1R1* were reduced in SnC EC2 cells, suggesting cell-type-specific alterations in stromal and vascular signaling programs (**Fig. 2b**). Notably, SnC EC2 cells were enriched for SASP factors with established roles in immune modulation and tissue remodeling, including *SPP1, GRN, LGALS9, TGFB1, MIF*, and *PDGFB*, highlighting the endothelial compartment as a source of signals relevant to endothelial-immune and endothelial-stromal interactions (**Fig. 2b** and **Supplementary Fig. 11a**).

To define the functional consequences of these transcriptional changes, we next performed gene set enrichment analysis (GSEA) using Hallmark and Reactome gene sets [45–47] (Supplementary Tables 10**–**13). GSEA identified significant enrichment (padj < 0.05) of pathways associated with ECM organization, cell adhesion, migration, cell cycle and apoptosis, vascular signaling, metabolism, inflammation, and additional tissue remodeling programs, with distinct enrichment patterns observed across SnC clusters (**Fig. 2c** and **Supplementary Figs. 11b, 12**). Upregulation of ECM-related gene sets was driven by *COL3A1*, *COL1A1*, *TNC, MMP11*, and *COL5A2* in SnC fibroblasts; *SPARC*, *COL4A1*, and *THY1* in SnC mural cells; and *SPARC*, *COL4A2*, *COL15A1*, and *SPP1* in SnC EC2 cells (**Supplementary Figs. 11b, 12**).

Enrichment of cell-cycle and apoptotic pathways reflected upregulation of genes including *NME1, VDAC1, UACA, BAX, CCND1, CCND2, CALR, YIF1A,* and *PDIA6* in SnC mural cells (**Supplementary Fig. 12a**). Similarly, enrichment of hypoxia-, inflammatory-, and vascular-associated programs in SnC EC2 cells was driven by upregulation of *ANGPTL4, IGFBP3, TGFB1*, and *SERPINA3* genes (**Supplementary Fig. 11b**). Together, these findings indicate that SnC states engage broad SASP programs that extend beyond matrix deposition to include inflammatory, vascular, and tissue-remodeling functions.

In addition to secreted factors (SASP), SnCs also exhibited upregulation of non-secreted cell-surface receptors (**Fig. 2d**). This was most evident in EC2 and mural populations, which upregulated collagen-binding integrins including *ITGA1* and *ITGB1*, forming the integrin α1β1 complex. SnC fibroblast cells exhibited elevated expression of receptors associated with non-canonical Wnt signaling, including *PTK7* and *ROR2*, which bind *WNT5A* and have been implicated in fibroblast activation (**Fig. 2d)**. In parallel, SnC EC2 cells upregulated receptors linked to vascular and immune signaling, including *UNC5B* and *TEK*, whereas SnC mural cells were distinguished by increased expression of *CD36*, a receptor implicated in the integration of vascular and metabolic cues (**Fig. 2d**). Together, these findings indicate that transcriptional programs differentiate SnC from non-SnC populations across cell types and functionally specialized SnC states through cell-type specific SASP and receptor programs. SnC fibroblast and mural populations predominantly expressed ECM-associated secretory programs that support matrix remodeling, whereas SnC EC2 cells were enriched for SASP factors and receptor programs associated with immune and vascular signaling within fibroids.

Consistent with the transcriptional expansion of EC2 cells in fibroids (**Fig. 1e,f**), these findings suggested a link between endothelial senescence and vascular remodeling, leading us further examine vascular structure and endothelial state.

### Senescent endothelial cells contribute to impaired vascular maturation in fibroids

Given the enrichment of SnC EC2 cells in fibroids and their SASP profiles associated with immune and stromal signaling, we next examined whether these transcriptional changes were reflected in vascular structure at the tissue level. CD31 IF staining demonstrated a reduction in vascular structures in fibroids relative to matched myometrium, accompanied by an increase in p16⁺ senescent cells (**Fig. 3a**). Quantitative analysis of Masson’s trichrome staining (MTS) further revealed a significant decrease in vessel lumen density in fibroids (*P* = 0.0011; **Fig. 3b** and **Extended Data Fig. 1a**), indicating impaired vascular organization in the context of ECM-dense fibroid tissue.

**Figure 3.**
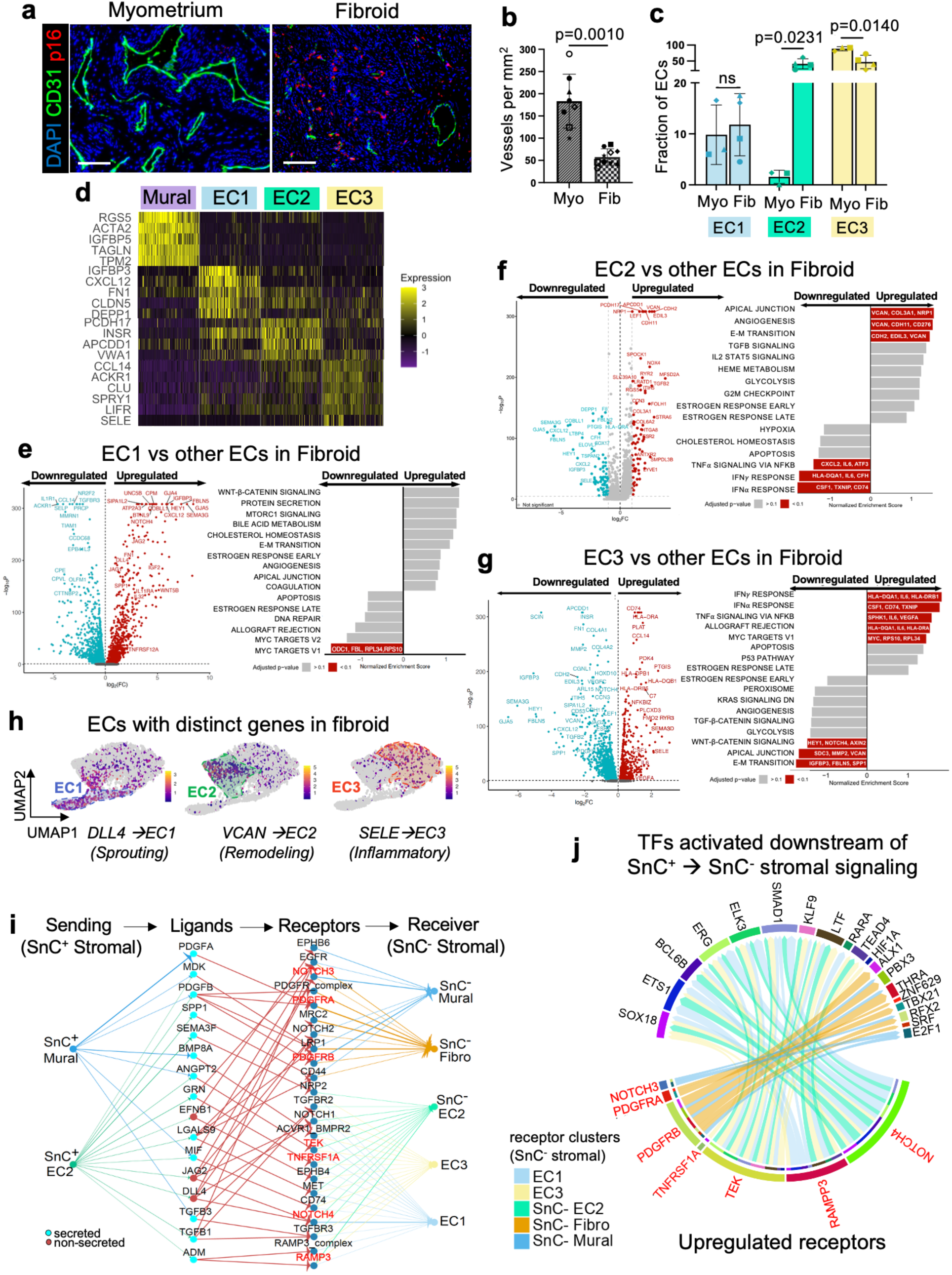
Senescence-associated endothelial remodeling contributes impaired vascular maturation in fibroids. **(a)** Representative immunofluorescence regions of interest (ROI) across a fibroid with matched myometrium sections stained for DAPI (blue, nuclei), CD31 (green, vasculature), p16 (red, senescence). **(b)** Quantification of vascular lumen density from MTS-stained sections showed a significant decrease in fibroids relative to matched myometrium. Statistical normality was assessed using the Shapiro–Wilk test, followed by a paired two-sided t-test (P = 0.0010). A total of 10 fibroid and 8 myometrium samples were analyzed, including 8 matched patient pairs used for paired statistical analysis. **(c)** Differential abundance of ECs between matched myometrium (Myo, *n* = 3) and fibroid (Fib, *n* = 4) samples in the scRNA-seq dataset. Bar plots show the relative fraction of EC1, EC2, and EC3 populations within ECs. Statistical significance was assessed using multiple paired two-tailed t-tests with Benjamini– Krieger–Yekutieli false discovery rate correction; adjusted *p* values are indicated.**(d)** Heatmap showing top DEG defining endothelial clusters (EC1, EC2, and EC3) and mural cells in fibroids. **(e–g)** Volcano plots showing differentially expressed genes in fibroid EC1, EC2, and EC3 populations relative to other fibroid endothelial populations (ECs). Upregulated genes are shown in red (log₂FC > 1, adjusted p-value < 10⁻⁵) and downregulated genes in blue (log₂FC < −1, adjusted p-value < 10⁻⁵), together with selected GSEA pathways. **(h)** Feature plots of key endothelial subcluster markers: *SELE* (inflammatory; EC3), *VCAN* (fibrotic/angiogenic; EC2), and *DLL4* (sprouting; EC1). **(i)** dominoSignal network plot showing signaling from SnC vascular cell ligands (mural and EC2 cells) to the receptors expressed in non-SnC stromal cell clusters (red circle, non-secreted; blue circle, secreted). The edge thickness from sending cluster to ligands and receptor to receiving cluster is proportional to average gene expression. **(j)** Chord diagram of dominoSignal analysis showing receptor-transcription factor (TF) correlations, linking receptors on non-SnC stromal cells, activated by ligands from SnC vascular cells, to downstream TF programs. The widths of the chords connecting receptors and TFs are proportional to receptor-TF correlation in the cluster.

Consistent with these structural observations, scRNA-seq analysis revealed a redistribution of endothelial populations in fibroids compared to matched myometrium. Although the overall proportion of endothelial cells was reduced in fibroids, clustering and differential abundance analysis identified a marked expansion of the EC2 population (**Fig. 3c**, **Supplementary Fig. 7**, and **Supplementary Table 3**). EC2 cells comprised approximately 1.5% of endothelial cells in myometrium but increased to ∼43% in fibroids, whereas EC3 cells, which predominated in myometrium, were proportionally reduced. Notably, ∼12% of EC2 cells were classified as SnC in fibroids. Similarly, ∼61% of mural cells were classified as SnC, representing the highest proportion of SnCs among all cell populations in fibroids, and indicating that SnCs are highly enriched within the vascular-associated compartment (**Figs. 1i** and **3c**; **Supplementary Table 3**).

To define functional differences among endothelial populations in fibroids, we performed differential gene expression and GSEA comparing each endothelial subcluster to the remaining endothelial cells (**Fig. 3d**–**g**, **Extended Data Fig. 1b** and **Supplementary Tables 14–20**). EC1 cells showed upregulation of vascular signaling-associated genes, including VEGF receptors (*FLT1, KDR*), *TEK*, and Notch pathway components (*DLL4, JAG2,* and *NOTCH4*), consistent with an activated endothelial signaling state (**Fig. 3e** and **Extended Data Fig. 1c**). Elevated expression of *DLL4*, *JAG2* and *NOTCH4* was consistent with Notch pathway engagement, whereas *PDGFB* expression suggested potential signaling to perivascular cells involved in vessel stabilization and maturation (Extended Data Fig. 1c).

In contrast, EC2 cells exhibited enrichment of genes associated with ECM interaction and remodeling, including *VCAN, COL3A1, COL6A2, CDH2, CDH11*, and *SPOCK1*, together with TGFβ-associated signaling components such as *TGFB2* (**Fig. 3f**). GSEA results further demonstrated enrichment of matrix and adhesion-related and angiogenesis pathways, whereas inflammatory programs, including TNFα signaling via NF-κB, and IFNγ/IFNα responses, were relatively suppressed. Although angiogenesis-related gene sets were enriched in EC2, this signal was primarily driven by ECM-associated genes (e.g., *VCAN, COL3A1*) rather than canonical VEGF-driven angiogenic programs (**Fig. 3f**). In addition, projection of the SenSig signature indicated that a subset of EC2 cells (∼14%) exhibited a senescence-associated transcriptional profile (**Fig. 1i** and **Supplementary Table 5**).

EC3 cells displayed an inflammatory endothelial phenotype characterized by increased expression of *SELE* (encodes E-selectin)*, SELP* (encodes P-selectin)*, VCAM1, IL6*, and *VEGFA*, consistent with endothelial activation and immune cell recruitment (**Fig. 3g**). GSEA results further showed enrichment of TNFα signaling via NF-κB, and IFNγ/IFNα responses, whereas pathways related to WNT/β-catenin signaling, apical junction organization, and epithelial– mesenchymal transition were relatively reduced (**Fig. 3g**).Feature-level visualization of *SELE, VCAN*, and *DLL4* further showed that inflammatory EC3 programs predominated in myometrium, whereas matrix-remodeling EC2 programs were expanded in fibroids, with sprouting EC1 signatures present in both tissues (**Fig. 3h** and **Extended Data Fig. 1d**).

Despite broad activation of angiogenesis-associated gene programs across endothelial populations, fibroids exhibited reduced vessel density (**Fig. 3a-b**), consistent with impaired vascular maturation. Given the expansion of SnC EC2 cells, we next examined SnC-associated alters stromal communication linked to vessel stabilization. We applied the ligand-receptor analysis with dominoSignal [48] to infer SnC EC2 cell signaling within the fibroid vasculature (**Fig. 3i** and **Supplementary Table 21**). DominoSignal, a cell-cell inference tool, infers intercellular ligand-receptor signaling based on gene expression and intra-cellular signaling by correlating receptor expression with TF activity scores. We focused on ligands differentially expressed in SnC EC2 population (adjusted p-value < 0.05, avg_log2FC > 0.3). SnC EC2 cells upregulated ligands such as *ANGPT2, JAG2*, and *DLL4* that engaged receptors on neighboring endothelial and mural populations, including *TEK* and *NOTCH* receptors (**Fig. 3i**).

*DLL4*- and *JAG2*-associated signaling through *NOTCH3* was maintained between endothelial and mural populations, consistent with preserved endothelial–mural communication involved in early vessel stabilization (**Fig. 3i**, **Extended Data Fig. 1e** and **Supplementary Fig. 13**). In parallel, SnC EC2-derived *PDGFB* signaling, which normally promotes pericyte recruitment during vessel maturation, was associated not only with PDGFRB interactions in mural populations but also with *PDGFRB*-, *PDGFRA*- and *LRP1*-associated receptor interactions in fibroblasts (**Fig. 3i** and **Extended Data Fig. 1e**). Downstream transcription factor analysis further showed that signaling from SnC EC2 and mural cells increased inferred activity of *PDGFRB*-associated transcriptional programs in fibroblasts, including *ALX1* and *PBX3* associated with mesenchymal activation [49, 50], and *TBX21* associated with inflammatory responses (**Fig. 3j** and **Supplementary Table 22**) [51]. By contrast, signaling from SnC EC2 and mural cells was associated with inferred *SOX18*, *ETS1*, *ERG*, and *SMAD1* activity downstream of *NOTCH4*-, *RAMP3*-, and *TEK*-associated signaling in ECs (**Fig. 3j**), consistent with adaptive vascular signaling responses. In addition, inferred *HIF1A* and *BCL6B* activity was enriched in ECs as a downstream result to ligands secreted by SnC mural and EC2 cells (**Fig. 3j**). Together, these findings suggest that senescence-associated vascular signaling in fibroids preserves features of canonical endothelial–mural communication while simultaneously engaging broader stromal signaling programs associated with fibroblast remodeling and inflammatory adaptation.

Although inflammatory EC3 cells were reduced in fibroids relative to myometrium, the fibroid-expanded EC2 population retained shared activation- and inflammation-associated markers including *SELP*, *CCL14*, and *TGFBR3*, suggesting persistence of endothelial inflammatory signaling despite endothelial compositional shifts (**Extended Data Fig. 1f**). This shift coincides with impaired vascular morphology and prompted further examination of how SnC stromal and vascular compartments engage immune populations in fibroids.

### Senescent stromal cells engage immune populations in fibroids

To determine whether SnC stromal cells are associated with immune engagement in fibroids, we first characterized immune cell populations by flow cytometry (**Supplementary Figs. 14** and **15; Supplementary Tables 23**–**25**). Analysis confirmed the presence of diverse immune populations in both fibroids and matched myometrium, including lymphoid populations (CD4⁺ and CD8⁺ T cells, γδ T cells, B cells, and natural killer (NK) cells) and myeloid populations (CD15⁺ granulocytes, CD14⁺HLA-DR⁺ macrophages, CD11c⁺HLA-DR⁺ dendritic cells, and CD16⁺ monocytes) (**Fig. 4a** and **Supplementary Fig. 15**). CD45⁺ cell counts normalized to tissue mass were reduced in fibroids compared to matched myometrium (**Supplementary Fig. 15a**), whereas the frequency of CD45⁺ immune cells remained comparable between the two tissues (**Supplementary Fig. 15b**). These flow cytometry analyses indicated an overall reduction immune cell count in fibroids, whereas IF staining showed redistribution of CD45⁺ cells throughout the fibroid tissue, compared to their predominant localization around lumen-like structures in myometrium (**Fig. 4b** and **Supplementary Fig. 15c**). Within the lymphoid compartment, the proportion of CD3⁺ T cells was significantly increased in fibroids relative to matched myometrium (**Fig. 4a**), while the relative distribution of CD4⁺, CD8⁺, and γδ T cell subsets remained largely unchanged (**Supplementary Fig. 15d**). IF staining for CD3 further demonstrated increased T cell infiltration throughout fibroid tissue (**Fig. 4b**). Within the myeloid compartment, granulocytes were significantly reduced in fibroids, whereas monocytes, macrophages, and dendritic cells were consistently detected without significant differences between fibroids and matched myometrium (**Fig. 4c** and **Supplementary Fig. 15c**). Notably, macrophages in both tissues were predominantly CD86⁺CD206⁻, consistent with a pro- inflammatory phenotype, with no significant shift in polarization state (**Supplementary Fig. 15e**).

**Figure 4.**
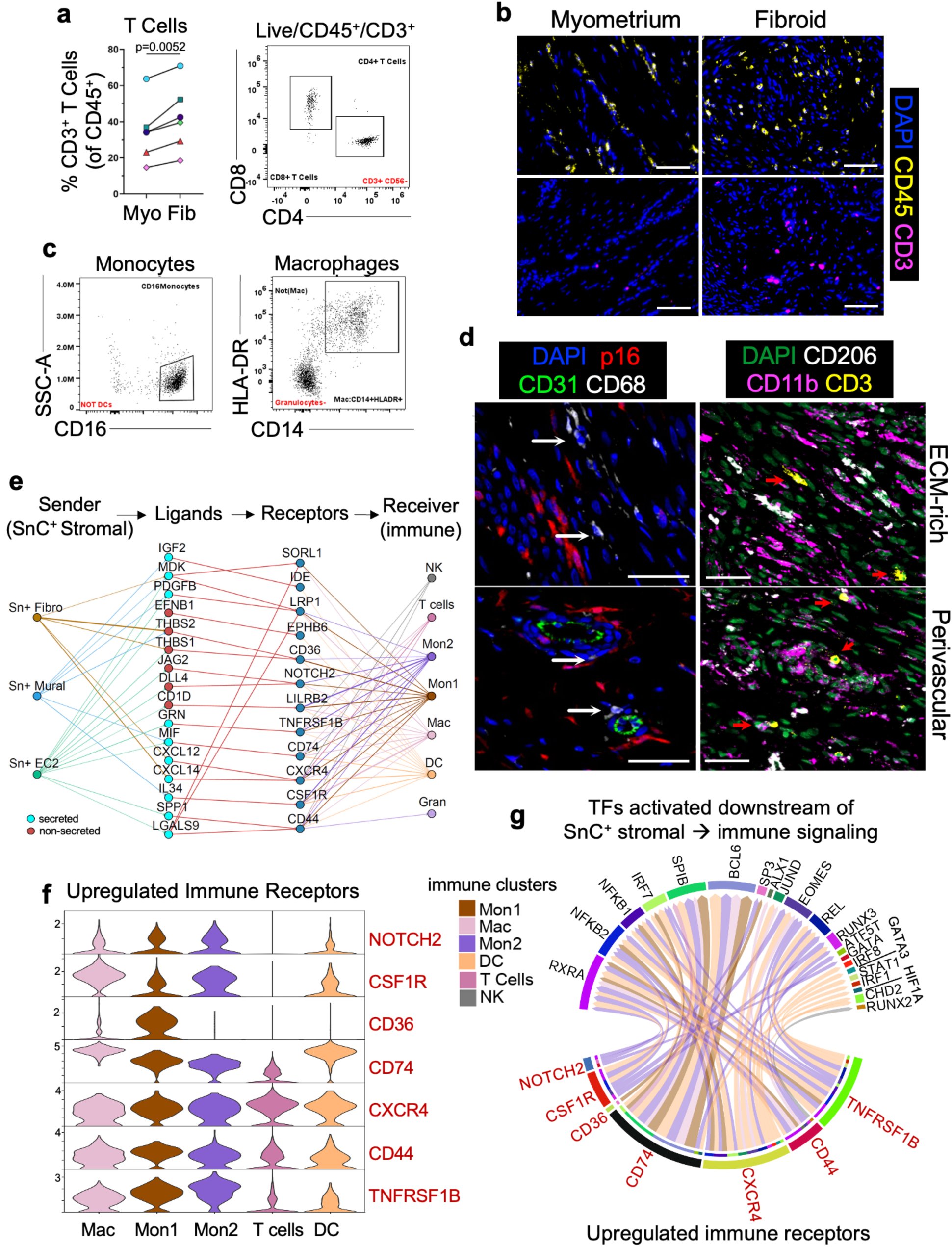
SnC stromal cells engage immune cells within fibroids. **(a)** The frequency of CD3+ T cells within fibroids and matched myometrium, quantified by flow cytometry, and the presence of CD8+ and CD4+ T cell subsets were observed (*P* = 0.0052, paired two-sided t-test; n = 6 matched samples). **(b)** Representative IF regions of interest across fibroid and matched myometrium sections stained with DAPI (blue, nuclei) together with either CD3 (magenta, T cells) or CD45 (yellow, leukocytes); scale bar = 100 μm. Representative IF regions of interest across a fibroid with matched myometrium sections stained for DAPI (blue, nuclei), CD3 (magenta, t cell), and CD45 (yellow, leukocytes), scale bar = 100μm. **(c)** The presence of populations of CD16+ monocytes, CD14+/HLA-DR+ macrophages in fibroid tissue were confirmed by using flow cytometry. **(d)** Representative immunofluorescence regions of interest showed two distinct spatial localizations of immune cells within fibroids. CD68⁺(white) myeloid cells are detected both adjacent to CD31⁺ (green) endothelial cells (perivascular niche) and within ECM-rich regions, where they mostly localize in close distance to p16⁺ SnC (red) cells. CD3⁺ T cells (yellow) showed a similar dual localization, whereas CD206⁺ myeloid cells (white) are enriched mostly in ECM-rich regions and CD11b⁺ myeloid (magenta) cells are observed in both perivascular and ECM-rich regions (Scale bar = 30μm). **(e)** DominoSignal network plot showing signaling from SnC+ stromal clusters to the immune clusters. The edge thickness from sending cluster to ligands and receptor to receiving cluster is proportional to average gene expression. **(f)** Violin plots showing the expression level of upregulated immune receptors (*CD36*, *CSF1R*, *CXCR4*, *TNFRSF1B*, *CD74*, *NOTCH2*, and *CD44*) across immune populations. **(g)** Chord diagram showing receptor-TF correlation linking immune receptor activation (as a results of ligand interaction sent by SnC stromal cells) to downstream TF activation within immune populations, showing SnC stromal cells activated TFs in myeloid cells rather than lymphocytes. The widths of the chords connecting receptors and TFs are proportional to receptor-TF correlation in the cluster.

To assess spatial relationships between immune cells and SnCs, we performed additional IF staining. CD68⁺ myeloid cells exhibited two primary localization patterns: one population adjacent to CD31⁺ vascular structures, consistent with a perivascular niche, and a second population distributed within ECM-dense stromal regions. In both contexts, CD68⁺ cells were frequently observed in proximity to p16⁺ cells, indicating a spatial association between myeloid cells and SnC stromal compartments (**Fig. 4d**). Similar spatial organization was observed for additional immune populations. CD3⁺ T cells were present in both perivascular and ECM-rich regions, paralleling the distribution of CD68⁺ cells. In contrast, CD206⁺ myeloid cells were primarily localized within ECM-dense regions, whereas CD11b⁺ myeloid cells were broadly distributed (**Fig. 4d**). Together, these spatial distributions indicate that immune cells localize to distinct SnC stromal compartments, including perivascular and ECM-rich niches, within fibroids.

scRNA-seq analysis further showed that myeloid cells in both tissues exhibited a shared inflammatory program marked by expression of *IL1B, CXCL8*, and *C5AR1* (**Extended Data Fig. 2a-b**). This signature was more prominent in myometrium but was reduced but persisted in fibroids, indicating ongoing myeloid inflammatory activity (**Extended Data Fig. 2b**). In contrast, T and NK cells in both tissues retained inflammatory and cytotoxic effector programs, although the relative expression of individual genes differed between tissues. *CCL5*, *IL32,* and *CCL4* showed relatively higher expression in fibroids. Conversely, *PRF1* and *GZMB* showed relatively higher expression in myometrium, while *GZMA* expression was similar between fibroids and matched myometrium (**Extended Data Fig. 2a,c,d**).

To examine potential SnC–immune interactions, we performed ligand–receptor analysis of the scRNA-seq dataset and examined signaling involving ligands that were differentially expressed in SnCs compared with the non-SnC fraction (adjusted *P* value < 0.05; average_log2FC > 0.3) (**Supplementary Table 26**). Our analysis revealed extensive predicted immune-directed signaling interactions originating from SnC stromal populations, particularly SnC EC2 cells (**Fig. 4e**). SnC EC2 cells showed increased expression of multiple SASP factors with established roles in immune recruitment and modulation, including *SPP1*, *GRN*, *LGALS9* and *MIF*, together with the Notch ligands *JAG2* and *DLL4*. These ligands were predicted to engage receptors such as *CD44*, *TNFRSF1B*, *CD74* and *NOTCH2* on myeloid and T-cell populations (**Fig. 4e**). SnC fibroblast and mural populations further contributed to predicted immune-associated signaling through ligand–receptor pairs, including *CXCL12*–*CXCR4*, *THBS1*/*THBS2*–*CD36* and *IL34*–*CSF1R* (**Fig. 4e**).

Immune populations within fibroids, particularly macrophage and monocyte subsets, expressed candidate receptors for SnC-derived signaling (**Fig. 4e**), including *CD36*, *CSF1R*, *TNFRSF1B*, *CD74* and *CXCR4* (**Fig. 4f**). Comparison with matched myometrium showed that these receptors were detectable across immune populations in both tissues, with cell-type-specific expression patterns (**Supplementary Fig. 16a**). IF staining of fibroid tissue showed abundant CD14⁺ myeloid cells co-expressing CD36, identifying a CD14⁺CD36⁺ myeloid population. CD3⁺ T cells were observed in close proximity to these CD14⁺CD36⁺ myeloid cells, indicating a spatial association and suggesting potential interactions between the two populations (**Supplementary Fig. 16b**). Together, these observations support a spatially organized immune microenvironment in fibroids, in which myeloid cells may respond to SnC-derived stromal cues and contribute to local immune modulation.

Transcription factor activity analysis identified inflammatory and immune-responsive programs involving *NF-κB*, *IRF* and *STAT* family factors (**Fig. 4g**). Predicted receptor–TF associations were concentrated in macrophage, monocyte and dendritic-cell populations, whereas T and NK cells showed comparatively limited receptor–TF coupling (**Fig. 4g** and **Supplementary Table 27**). In myeloid clusters, receptors including *CD74*, *CSF1R*, *CD36*, *CXCR4* and *TNFRSF1B* were associated with multiple TF programs, consistent with activation of inflammatory and immune-responsive pathways. DominoSignal analysis showed that *MIF* derived from SnC EC2/mural populations engages its receptor *CD74* in myeloid cells, that was strongly associated with activation of *BCL6* and *SPIB* TFs (**Fig. 4g**). Together, these findings suggest that myeloid cells represent the principal immune compartment transcriptionally responsive to SnC-derived stromal signals.

After analysis of signaling from SnC stromal populations to immune cells, we next assessed the reverse direction (**Supplementary Fig. 16**). Immune-to-SnC EC2 interactions showed a distinct pattern. Myeloid-derived *VEGFA* was linked to *FLT1, KDR*, and *NRP1* on EC2 cells, together with *ADM-RAMP3* signaling, consistent with vascular-related responses. In addition, EC2 cells expressed *ACKR1*, which binds to and presents chemokines produced by myeloid (*CXCL8, CXCL5*) and lymphoid (*CCL5*) cells (**Supplementary Fig. 16c**). In contrast, immune-to-SnC interactions were largely similar for SnC mural and fibroblast populations.

Immune-derived *LGALS9* showed interaction with *LRP1* and *MRC2*, and PDGF ligands (*PDGFB, PDGFC,* and *PDGFD*) were associated with *PDGFR* receptors on both SnC mural and fibroblast cells, indicating shared stromal activation pathways (**Supplementary Fig. 16d**,**e**).

Together, these findings indicate that SnC stromal cells actively engage immune populations in fibroids and contribute to a coordinated vascular-stromal-immune microenvironment.

### Immune-associated mechanosensing programs are linked to SnC stromal niches in fibroids

Histological analysis using MTS demonstrated a marked increase in collagen-rich ECM regions in fibroids, accompanied by a reduction in smooth muscle-dominated areas compared with matched myometrium (**Fig. 5a** and **Extended Data Fig. 1a**). Consistent with this ECM-dense architecture, unconfined compression testing revealed an approximately twofold increase in Young’s modulus in fibroid tissue, indicating increased tissue stiffness (**Fig. 5b**). Three-dimensional CODA imaging of a representative matched pair further showed dense ECM organization and reduced vascular representation in fibroid tissue relative to myometrium (**Fig. 5c** and **Supplementary Fig. 17a**). Together, these findings established matrix accumulation and increased stiffness as defining features of the fibroid microenvironment.

**Figure 5.**
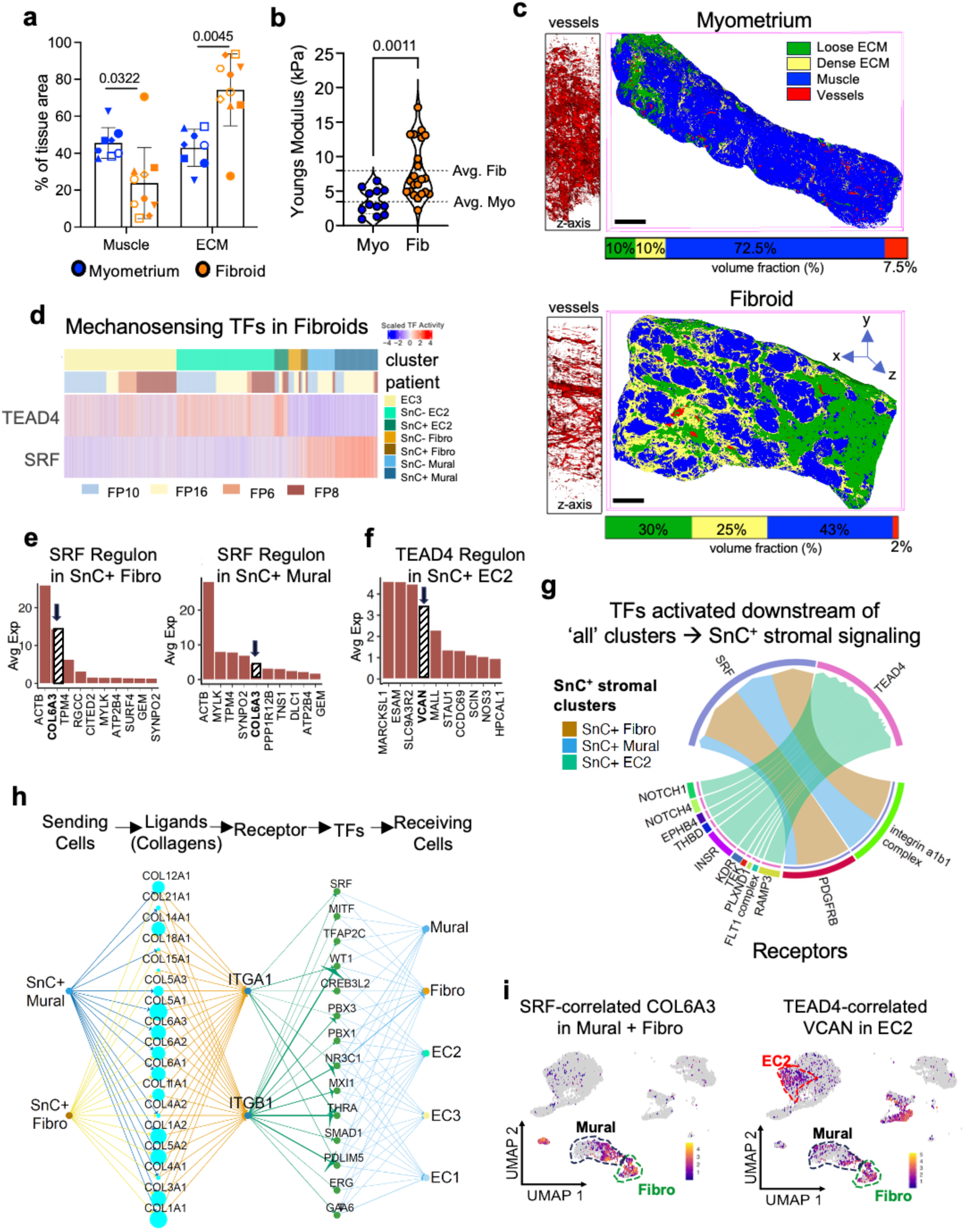
Immune- and stromal-associated mechanosensing programs are linked to SnC-associated ECM remodeling in fibroids. (**a**) QuPath-based segmentation of MTS-stained sections showing reduced muscle area and increased ECM area in fibroids (*n* = 10 patient samples) compared with matched myometrium (*n* = 8 patient samples). Eight matched fibroid–myometrium pairs were included in the paired statistical analyses; the two unmatched fibroid samples are shown but were not included in the statistical comparisons. Normality of the paired differences was assessed using the Shapiro–Wilk test, followed by a two-sided paired *t*-test (muscle, *P* = 0.0322; ECM, *P* = 0.0045). Data are presented as mean ± s.d. Individual points represent patient samples, and identical symbols indicate matched pairs. (**b**) Unconfined compression testing showing an approximately twofold increase in Young’s modulus in fibroid tissue, indicating increased tissue stiffness. A total of 22 fibroid and 12 myometrium tissue specimens were analyzed. Normality was assessed using the Shapiro–Wilk test, followed by a two-sided unpaired *t*-test (*P* = 0.0011). Individual measurements are shown, and dashed lines indicate group means. (**c**) Representative three-dimensional CODA reconstruction of matched myometrium and fibroid tissues showing loose ECM (green), dense ECM (yellow), vessels (red) and muscle (blue). The relative reconstructed volume fraction of each segmented compartment is shown below the corresponding reconstruction. Orthogonal vessel views are shown at left; x–y denotes the section plane and z denotes serial-section depth. Scale bar, 2 mm. (**d**) Heatmap of scaled SCENIC regulon activity showing higher *TEAD4* regulon activity in SnC EC2 cells and higher *SRF* regulon activity in SnC fibroblast and mural populations relative to their non-SnC counterparts across patients. (**e**) Top ten *SRF* regulon target genes ranked by mean expression in SnC fibroblast and mural cells. (**f**) Top ten *TEAD4* regulon target genes ranked by mean expression in SnC EC2 cells. (**g**) Chord diagram showing receptor–TF correlations associated with *SRF* and *TEAD4* regulon activity across SnC stromal populations. Chord width is proportional to the within-cluster receptor–TF correlation. Associations with a data-wide Spearman correlation > 0.25 and a within-cluster Spearman correlation > 0.12 are shown. (**h**) dominoSignal network showing predicted collagen signaling from SnC fibroblast and mural cells through integrin α1β1-containing receptors and associated TF programs in stromal receiving populations. Edge thickness from sending populations to ligands is proportional to average ligand expression, edge thickness from receptors to TFs is proportional to the data-wide receptor–TF correlation, and edge thickness from TFs to receiving populations is proportional to median TF activity. (**i**) UMAP feature plots showing *VCAN* and *COL6A3* expression across fibroid endothelial and stromal populations.

Given these changes in tissue mechanics, we next examined whether mechanosensing transcriptional programs were enriched in SnC stromal populations (fibroblast, mural and EC2) relative to their non-SnC counterparts. We used SCENIC to infer TF regulon activity from the scRNA-seq data and compared regulon activity scores between SnC and non-SnC cells using a Wilcoxon test (**Supplementary Table 28**). Among the differentially active regulons, we focused on the mechanosensing-associated regulators *SRF* and *TEAD4*. *SRF* regulon activity was increased in SnC fibroblast and mural populations, whereas *TEAD4* regulon activity was increased in SnC EC2 cells relative to their non-SnC counterparts (FDR < 0.05; effect size > 0.2) (**Fig. 5d** and **Supplementary Fig. 17b**,**c**).

We further examined the TF target genes within the *SRF* and *TEAD4* regulons identified by SCENIC (**Supplementary Table 29**). In SnC fibroblast and mural populations, the ten *SRF* regulon genes with the highest average expression included cytoskeletal and contractile genes, including *ACTA2* and *MYLK*, together with ECM components, most prominently *COL6A3* (**Fig. 5e**). In contrast, SnC EC2 cells exhibited a distinct mechanosensing program characterized by *TEAD4* regulon activity. The genes with the highest average expression within the *TEAD4* regulon included endothelial-remodeling genes, including *VCAN* (**Fig. 5f**).

To define upstream signaling inputs, we again used dominoSignal for ligand–receptor analysis. Expression of *PDGFRB* and *integrin α1β1* receptor components was correlated with SRF regulon activity in SnC fibroblast and mural populations, whereas *RAMP3* and *INSR* expression was correlated with *TEAD4* regulon activity in SnC EC2 cells (**Fig. 5g** and **Supplementary Table 30**). Examination of the associated ligands indicated that *PDGFB* and *PDGFD* were predicted to signal through *PDGFRB* in SnC fibroblast and mural cells, whereas immune-derived *ADM* was predicted to signal through *RAMP3* in SnC EC2 cells (**Supplementary Figs. 18**, **19**). These findings suggest that immune and stromal signaling inputs converge on mechanosensitive programs within SnC populations.

Network analysis further connected collagen ligands, including *COL6A3*, from SnC fibroblast and mural populations to *integrin α1β1* and associated TF programs, consistent with a potential feed-forward program that may reinforce matrix deposition (**Fig. 5h** and **Supplementary Table 31**). Feature-level visualization showed that *COL6A3* expression was concentrated in fibroblast and mural populations, whereas *VCAN* expression was predominantly localized to EC2 cells among fibroid ECs (**Fig. 5i**), consistent with cluster-specific matrix-associated outputs of mechanosensing programs.

Integration of ligand–receptor and TF analyses with dominoSignal indicated that immune- and stromal-derived signals were associated with mechanosensitive programs within SnC stromal populations. *PDGFRB*-associated signaling in fibroblast and mural cells corresponded with SRF regulon activity and *COL6A3* expression, whereas *ADM*–RAMP3 signaling in SnC EC2 cells was associated with *TEAD4* regulon activity and *VCAN* expression (**Fig. 5g** and **Supplementary Fig. 17**). These observations are consistent with a model in which immune-derived cues contribute to mechanosensitive transcriptional programs across stromal compartments. Overall, ECM stiffening in fibroids was associated with distinct mechanosensing programs in SnC stromal populations, with *SRF*-associated programs in fibroblast and mural cells linked to matrix deposition and *TEAD4*-associated programs in EC2 cells linked to endothelial remodeling. Together, these findings link ECM remodeling and SnC-associated mechanosensitive TF programs with immune- and stromal-derived signaling inputs within fibroids.

### Matrix degradation is associated with reduced senescent cell burden in human fibroids

To examine the relationship between ECM organization and SnC states in fibroids, we analyzed p16⁺ cells in fibroid tissues obtained from a clinical collagenase injection study [52]. Collagenase-based therapies have been clinically used to degrade pathological collagen-rich matrices, most notably in Dupuytren’s contracture [53] and Peyronie’s disease [54–56], where enzymatic disruption of collagen improves tissue remodeling and function. More recently, localized collagenase injection has been explored as a minimally invasive approach for treating uterine fibroids, targeting the dense ECM that characterizes these tumors and contributes to their stiffness and persistence [52, 57]. In that study, fibroids were locally injected with saline or collagenase (0.1 or 0.2 mg per cm³ tissue volume) 60-90 days prior to surgical resection [52] (**Fig. 6a**).

**Figure 6.**
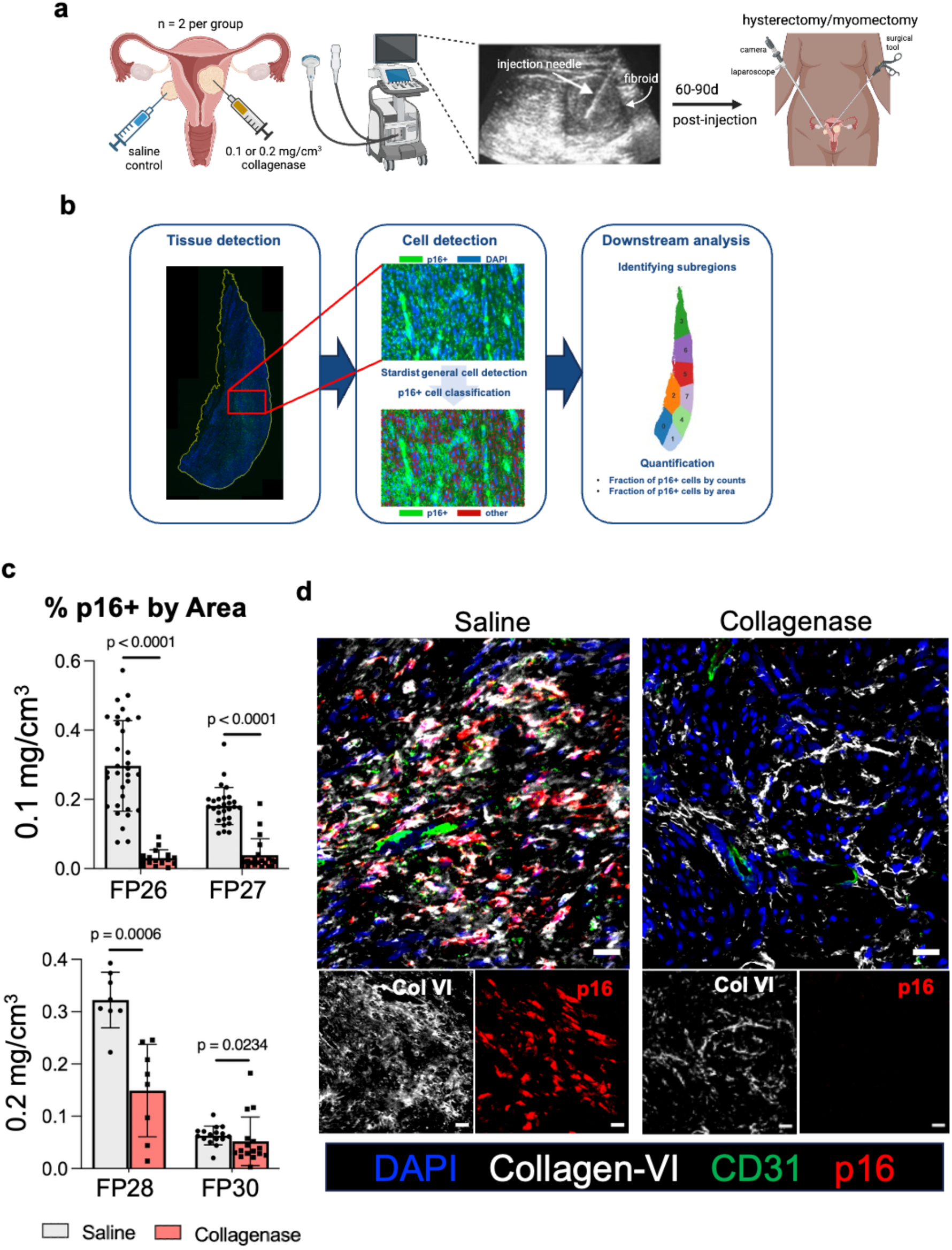
Collagenase-mediated matrix degradation is associated with reduced p16⁺ SnC burden in uterine fibroids. **(a)** Schematic overview of ultrasound-guided intrafibroid injection of collagenase (0.1 or 0.2 mg/cm³) or saline control, followed by hysterectomy or myomectomy 60–90 d after injection. Two patients were included in each collagenase dose group. **(b)** Automated image-analysis workflow comprising whole-tissue detection, StarDist-based cell detection, classification of p16⁺ cells, spatial partitioning of tissue into subregions and quantification of p16 positivity using cell count- and area-based measurements. **(c)** Area-based p16⁺ fraction within individual tissue subregions of saline- and collagenase-treated fibroids from patients receiving 0.1 mg/cm³ collagenase (FP26 and FP27) or 0.2 mg/cm³ collagenase (FP28 and FP30). Points represent individual tissue subregions; bars indicate the median and error bars indicate the interquartile range. Normality of the subregion-level values was assessed separately for each patient using the Shapiro–Wilk test. Saline- and collagenase-treated subregion distributions were compared within each patient using the Mann–Whitney U test. Exact *P* values are shown in the figure. **(d)** Representative IF regions of interest (ROIs) from fibroid sections of patient FP29 showing DAPI (blue), collagen VI (white), p16 (red), and CD31 (green) following high-dose collagenase treatment (0.2 mg/cm³) or saline control. Scale bar, 20 μm.

Histological analysis showed that saline-treated fibroids exhibited dense tissue architecture with p16⁺ cells distributed throughout the analyzed tissue regions (**Supplementary Fig. 20a-c**). In contrast, collagenase-treated fibroids displayed reduced tissue density and structural gaps consistent with matrix degradation (**Supplementary Fig. 20a**,**c**). To account for these architectural differences, p16⁺ cell burden was quantified using both cell count-based and area-based metrics (**Fig. 6b–c**, **Supplementary Fig. 20b** and **Supplementary Table 32**).

Quantitative analysis demonstrated a reduction in p16⁺ cell burden in collagenase-treated fibroids compared to saline-treated controls at both the 0.1 and 0.2 mg cm⁻³ doses (**Fig. 6c** and **Supplementary Fig. 20b**). Subregion-level analysis nevertheless revealed marked spatial heterogeneity in p16⁺ cell distribution, with p16⁺ cells concentrated within discrete regions of the fibroid rather than uniformly distributed throughout the tissue. Although collagenase treatment was associated with lower overall p16⁺ cell abundance, variation among tissue subregions remained evident across several samples (**Fig. 6c**), suggesting spatially localized accumulation of p16⁺ SnC cells within the fibroid microenvironment.

To further examine the relationship between ECM organization and p16⁺ cell localization, we performed IF staining for collagen VI, CD31 and p16 (**Fig. 6d** and **Supplementary Fig. 20c**). Collagenase-treated samples showed reduced collagen VI signal and fewer p16⁺ cells relative to saline-treated controls. In saline-treated tissues, p16⁺ cells were observed within collagen VI-rich regions and near CD31⁺ vascular structures (**Fig. 6d** and **Supplementary Fig. 20c**). The accumulation of collagen VI in control tissues was consistent with the SRF-associated ECM program and increased *COL6A3* expression identified in SnC stromal populations (**Fig. 5**).

In sum, these results support a relationship between ECM organization and SnC in fibroids, in which collagen-rich matrix environments are associated with increased p16^+^ SnC accumulation, whereas collagenase-mediated matrix degradation is associated with reduced p16^+^ SnC burden. In the context of the mechanosensing and immune-associated signaling programs described above, these findings are consistent with a model in which ECM structure contributes to the maintenance of SnC-associated stromal states within the fibroid microenvironment.

## Discussion

Uterine fibroids are characterized by dense ECM deposition, altered vascular structure, and progressive tissue stiffening [2, 7], but the cellular programs coordinating these features remain incompletely defined. In this study, we identify SnC states as a central organizing feature of the fibroid microenvironment, distributed across stromal and vascular compartments and associated with coordinated changes in matrix remodeling, vascular signaling, immune engagement, and mechanosensing pathways. Although the number of patient samples analyzed for scRNA sequencing (*n* = 4 fibroid and *n* = 3 matched myometrium samples) is modest and may limit the full capture of fibroid heterogeneity, the convergence of spatial immunofluorescence, transcriptional, and histological data across an extended cohort (*n* = 30 patients) supports the robustness of these observations.

Rather than representing a uniform state, senescence in fibroids is heterogeneous and compartmentalized, with distinct SnC populations (“senotypes”) identified in fibroblast, mural, and endothelial compartments. These senescent states exhibit specialized transcriptional programs and signaling outputs, consistent with the concept that senescence is a context-dependent program shaped by cell identity and tissue environment [17, 58, 59]. Notably, SnC fibroblast and mural populations are enriched for ECM-associated SASP factors, whereas SnC ECs exhibit SASP profiles linked to immune modulation and vascular signaling, indicating functional specialization across SnC compartments. Interpretation of these patterns should be considered in the context of matched myometrium controls, which, although patient-matched, may not fully represent normal uterine tissue.

Our data further suggest that SnC ECs are associated with altered vascular organization in fibroids. Despite enrichment of angiogenesis-related gene programs, fibroids exhibit reduced vessel density and impaired lumen formation. This apparent disconnect between angiogenic signaling and vascular structure is consistent with prior observations that fibroids are supplied by peripheral arterial plexuses yet maintain hypovascular cores [60]. These findings raise the possibility that blood supply alone is insufficient to explain fibroid maintenance or therapeutic responses such as uterine artery embolization [61]. Rather, fibroid persistence may reflect adaptation to a hypoxic, ECM-dominated microenvironment, supported by coordinated stromal and vascular signaling programs. However, the extent to which these processes function independently of vascular supply cannot be determined from the current data.

In parallel, we identify extensive immune engagement within senescent stromal compartments, supported by both spatial and transcriptional analyses. Immune cells localize to perivascular and ECM-rich regions and are frequently positioned adjacent to SnC. Senescent stromal populations express SASP factors with established roles in immune recruitment and modulation, and immune populations exhibit receptor expression and transcriptional programs consistent with responsiveness to these signals. These findings suggest that SnC participate in coordinated vascular-stromal-immune interactions, contributing to the establishment of complex microenvironmental niches within fibroids. It should be noted that inference of these intercellular signaling interactions is based primarily on a single computational framework (dominoSignal) and has not been independently validated, and therefore represents putative interactions requiring further confirmation. Notably, several of the inferred senescence signaling interactions are consistent with pathways previously implicated in fibroid pathogenesis, including TGFβ- [62], PDGF- [63] [64], and Wnt/β-catenin [65], providing additional biological context for these observations.

A defining feature of fibroids is increased matrix stiffness, and our data link this mechanical phenotype to senescent cell states. Senescent stromal populations are predicted to activate mechanosensitive transcription factors, including *SRF* in fibroblast and mural cells and *TEAD4* in endothelial cells, with downstream programs associated with cytoskeletal organization and ECM production. Integration of ligand-receptor and dominoSignal analyses suggests that immune- and stromal-derived signals converge on these mechanosensitive pathways. These observations are consistent with a model in which mechanosensing, ECM remodeling, and senescence are coupled, forming a system that sustains fibrotic tissue organization. However, because these findings are derived from human tissue samples, the ability to perform controlled perturbation experiments to establish causality is limited, particularly in the absence of a tractable in vivo model of fibroids.

Hormonal signaling represents an additional consideration. Estrogen and progesterone are key drivers of fibroid growth, and their receptors (ESR1 and PGR) are primarily expressed in smooth muscle cells (SMCs) [66]. Due to limited recovery of SMCs in our scRNA-seq dataset, likely reflecting known size-dependent biases and technical challenges associated with microfluidic capture [67], we were unable to robustly assess the relationship between senescence and hormone receptor expression within this population. Although recovered SMCs recapitulated expected increases in ESR1 and PGR expression in fibroids, they were insufficient for reliable comparison of senescent and non-senescent states, leaving open questions regarding the interaction between hormonal signaling and senescence in fibroid biology.

Importantly, analysis of human fibroids following collagenase treatment provides translational support for the relationship between ECM organization and senescent cell states. Collagenase, which has established clinical use in collagen-rich fibrotic conditions, reduces ECM density and is associated here with a decrease in SnC burden. These findings suggest that matrix organization may contribute to the maintenance of SnC states. However, collagenase is not selective for pathological ECM and may disrupt the structural integrity of adjacent normal tissue. Given the mechanically dynamic nature of the uterus, excessive matrix degradation could impair myometrial function, and matrix remodeling may transiently generate bioactive matrikine fragments that influence immune or stromal signaling [68]. Thus, while these data support a relationship between ECM structure and senescence, targeted modulation of mechanosensitive pathways may represent a more precise therapeutic strategy than broad enzymatic ECM degradation.

Together, this work provides a framework for understanding fibroids as senescence-associated tissue systems, in which heterogeneous SnC states are linked to ECM remodeling, vascular dysfunction, immune engagement, and mechanosensitive signaling. These findings extend the role of senescence beyond a static marker of aging to an active component of tissue organization and disease progression. More broadly, this study suggests that targeting SnC states or their associated microenvironmental interactions, including ECM remodeling and mechanosensing, may represent a therapeutic strategy for fibroids and other fibrotic diseases.

## Materials and Methods

### Tissue sample collection

This study was approved by the Johns Hopkins University Institutional Review Board (IRB00359032 and IRB00091412 and IRB00196175). Fibroid and myometrium tissue were collected from consented patients undergoing elective hysterectomy at Johns Hopkins Hospital. Patient demographics, clinical characteristics, and sample allocation across experiments are summarized in **Supplementary Table 1**. Prior to patient surgery, a screening ultrasound was performed and after surgery, a histopathologic exam was performed to confirm diagnosis of uterine fibroids. Samples were collected from the operating room and processed under sterile conditions and immediately transported to the lab in either formalin or Hank’s balanced salt solution (HBSS) at 4°C.

### Tissue digestion optimization

Fresh fibroid tissue (1 g) obtained from a single patient was used to evaluate the impact of dissociation conditions on cell yield and cellular heterogeneity. Four dissociation protocols were tested: (i) standard manual dissociation using 3 mg/mL Type I collagenase (Worthington Biochemical Corporation, LS004197), (ii) a custom gentleMACS Octo Dissociator program (Miltenyi Biotec) using 1 mg/mL collagenase supplemented with Enzyme R (Miltenyi Biotec; 10 µL or 20 µL), and (iii-iv) predefined gentleMACS programs optimized for soft and hard tumor dissociation (Miltenyi Biotec) using Enzyme R (10 µL or 20 µL). All digestions were performed in RPMI 1640 medium (Gibco) supplemented with HEPES (Gibco) and DNase I (1 mg/mL; Sigma-Aldrich). Following enzymatic digestion, samples were filtered through a 100 µm cell strainer (Corning), washed with phosphate-buffered saline (PBS; Gibco), and centrifuged at 500 × g for 5 min. Cell pellets were resuspended in PBS and passed through a 40 µm cell strainer (Corning, CLS431750) to generate single-cell suspensions. Cells were stained with antibodies (see **Supplementary Table 23** for antibody details) and analyzed using an Aurora spectral flow cytometer (Cytek Biosciences). Flow cytometry acquisition and gating strategies are described in the Flow Cytometry section. Based on these optimization experiments, the standard manual dissociation protocol was used for all subsequent flow cytometry and scRNA-seq experiments.

### Tissue dissociation

Tissues were finely minced and digested in RPMI 1640 medium (Gibco) with 3 mg/mL type I collagenase (Worthington Biochemical Corporation) and 1 mg/mL deoxyribonuclease I (DNase I) from bovine pancreas (Sigma-Aldrich) for 45 min at 37 °C. Digested tissues were ground through a 100 µm cell strainer (Miltenyi Biotec) with excess RPMI and 1 x PBS. The cell suspension was centrifuged, then washed again with 1 x PBS and passed through a 40 cell µm strainer (Miltenyi Biotec).

### Flow cytometry

Single-cell suspensions from each fibroid and matched myometrium were divided equally between two antibody panels: Myeloid (**Supplementary Tables 23** and **24**) and Lymphoid (**Supplementary Table 25**), which were processed and stained separately. Both panels were stained in 100 µL of Zombie NIR Fixable Viability Dye (BioLegend) at a 1:7000 dilution for 30 min at 4°C, then washed twice with BD wash buffer (BD Biosciences). For the Lymphoid panel, staining was performed at room temperature and in two steps; the samples were first stained for 15 minutes with γδ TCR antibody (diluted with Fc block (BioLegend), Monocyte block (BioLegend), and BD wash buffer). Then the rest of the surface antibodies were added to the samples as a concentrated stain cocktail (prepared in Super Bright buffer (ThermoFisher Scientific)) for a final stain volume of 100µL. Samples were incubated at room temperature for an additional 30 minutes. For the antibody panels, a concentrated antibody cocktail was prepared in Monocyte block and Super Bright Buffer then diluted with BD wash buffer just prior to staining (**Supplementary Table 33**). Samples were incubated in 100 µL of the Myeloid antibody cocktail for 45 minutes at 4°C. After staining, both panels were fixed with 100 µL of FluoroFix buffer (BioLegend) for 15 min at room temperature, washed twice, and resuspended in BD Wash Buffer, then stored overnight at 4 °C. Samples were analyzed the following day on an Aurora Spectral Flow Cytometer (Cytek Biosciences).

### Western blotting

Freshly isolated myometrial (*n* = 9) and fibroid tissues (*n* = 9) from patient-matched tissues were finely chopped and incubated overnight at 37 °C with constant agitation in Hanks’ Balanced Salt Solution (HBSS; Gibco, cat. no. 14025-092) supplemented with 1.5 mg/ml type

IV collagenase (Gibco, cat. no. 17104-019) or type I collagenase (Sigma, cat. no. C0130-1G), DNase I (Sigma, cat. no. D5025-150KU), 2 mM CaCl₂ (Quality Biological, cat. no. 351-130-721), 10 mM HEPES (Gibco, cat. no. 15630-080), gentamicin (Sigma, cat. no. G1397), and normocin (InvivoGen, cat. no. ant-nr-1). Following enzymatic digestion, the resulting cell suspension was filtered through 100 μm cell strainers (CELLTREAT, cat. no. 229485) and centrifuged twice at 1500 rpm for 10 minutes. The supernatant was discarded, and the resulting cell pellets were lysed in RIPA buffer (Sigma, R0278) supplemented with a protease and phosphatase inhibitor cocktail (Cell Signaling Technology, 5872S) to prepare protein lysates for western blot analysis. The Pierce™ BCA Protein Assay Kit (ThermoFisher Scientific, 23227) was used to quantify protein concentrations. To perform electrophoresis, 30 µg of protein was loaded onto 4–12% NuPAGE gels (Invitrogen, NP0321BOX) and separated under reducing conditions by SDS-PAGE. Proteins were then transferred to nitrocellulose membranes using the iBlot 3 transfer stack (Invitrogen, IB33002) and the iBlot 3 Western Blot Transfer System (Invitrogen, IB31001). The transfer efficiency was verified with Ponceau S solution (Sigma, P7170). Membranes were cut based on the molecular weights of the target proteins and blocked for 20 min in 5% non-fat milk (BIO-RAD, 1706404) diluted in TBST (Tris-Buffered Saline with 0.1% Tween 20). Primary antibodies specific to p16 (Abclonal, A11651) and β-actin (Cell Signaling Technology, 4970S) were applied for 1 h at room temperature. Following three washes with TBST (5 min each), HRP-conjugated secondary antibody (1:5000 dilution; GE Healthcare, NA934V) was added and incubated for 1 hour at room temperature in 5% non-fat milk with TBST. After another set of three 5-minute washes, the immunoreactive bands were visualized using the SuperSignal™ West Atto Ultimate Sensitivity Chemiluminescent Substrate (ThermoFisher Scientific, A38556) and captured using an Azure Imager c300 system (Azure Biosystems). Band intensities were quantified with ImageJ 1.52a software, normalized to the loading control β-actin.

### scRNA-seq sample preparation

For scRNA-seq the cells were ACK Lysis buffer then run through a MACS column using the dead cell removal bead kit (Miltenyi Biotec) following the manufacturer’s protocol. CD45 MicroBeads (Miltenyi Biotec) were then used to select the CD45^+^ cells. These were then mixed 1:1 with the CD45^-^ flow-through fraction to generate the final CD45-enriched cell suspension.

### scRNA-seq analysis

Samples were aligned using Cell Ranger (v6.1.1) to the GRCh38 2020-A reference [69]. High-quality cells were selected by removing cells with fewer than 500 UMI, fewer than 350 unique features, or greater than 25% mitochondrial gene expression. Stress-related genes were removed for PCA and clustering, though they were included in differential expression. UMI count was regressed during data scaling. The data were integrated by patient using Harmony with default settings and 30 PCs. The first 30 corrected PCs were used for Louvain clustering with a resolution of 0.3. Differential expression in a given cluster compared to remaining cells was performed using Wilcoxon-rank sum test with false discovery rate (FDR) multiple testing correction. Ranked gene set enrichment analysis was performed using the fgsea package and Hallmark gene sets from MSigDB [46, 70, 71].

The R package edgeR [34] was used to test for changes in cell type abundance between tissues as described in Orchestrating single-cell analysis with Bioconductor book [72]. The scRNA-seq data was summarized to obtain the number of cells in each cell type for each patient and tissue. The patient identity was included as a covariate in the design matrix and edgeR’s glmQLFTest was used to test for differences in cell type abundance.

### Transfer learning of the senescence signature and analysis of senescent cells

The transfer learning tool projectR (version 1.22.0) [39, 40] was used in combination with the *in vivo*-derived senescence signature, SenSig [38], to label putative SnCs. Briefly, given the SenSig, projectR was used to perform a generalized least-squares fit to the z-scored log-normalized counts from the single-cell dataset. The cells with positive coefficients from the projectR analysis (projection weights) and p-values below 0.01 were labeled as senescent cells. The senescent cells were then compared to the non-senescent fraction within the Fibroblast, Mural, and EC2 clusters using the findMarkers function in Seurat (version 5.2.1) [73] with default parameters to obtain a list of differentially expressed genes. The CellPhoneDB database (version 4.0.0) [74] was used to annotate signaling genes that are also secreted as putative SASP. MatrisomeDB (2.0) [42] was used to annotate ECM genes, and “The in silico human surfaceome [75]” publication by Bausch-Fluck et al. was used to annotate the genes encoding cell-surface proteins. We then used the R package fgsea (1.32.4) [45] with the Hallmark and Reactome gene sets (MSigDB, version 10.0.2) [46, 70, 71] to find pathways that are enriched in the senescent cells compared with the non-senescent populations. To identify the projection drivers, we used the differentially expressed genes from the FindMarkers results with FWER-corrected p-value < 0.05 and average log2 fold change > 0.3 and subset to the genes upregulated in SenSig. We additionally used the projectionDriveR [41] function in the projectR package to obtain confidence interval bound differences in mean expression between the SnC and non-SnC populations for the genes upregulated in SenSig.

### Cell-cell signaling analysis with dominoSignal

The R package dominoSignal (version 0.2.2) [76] was used to perform cell-cell signaling analysis. The CellPhoneDB database (version 4.0.0) [74] was used as the interaction database for the dominoSignal analysis. The cells were first scored for transcription factor activity using the Python package pySCENIC v0.12.1 [77, 78] and dominoSignal was used to infer signaling networks in the data, prioritizing interactions where the receptor expression was correlated with transcription factor activity. dominoSignal was run using the parameters use_clusters = T, use_complexes = T, remove_rec_dropout = F, min_tf_pval = .001, max_tf_per_clust = Inf, max_rec_per_tf = Inf, rec_tf_cor_threshold = .25. This analysis was performed per patient sample, and interactions found in at least 3 fibroid samples were considered further. To infer signaling from SnCs, we additionally filtered the ligands to include genes that were differentially expressed in SnCs compared with non-SnCs. To infer TF-receptor associations, we assessed correlation of receptor expression and TF activity within clusters. TF activity scores in the SnC cells were compared to the non-SnC fraction using Wilcoxon test and calculated effect sizes using rstatix (version 0.7.3). The R package igraph (version 2.1.4) was used to generate the network plots.

### Compression testing of fibroid and matched myometrium tissues

Fresh uterine fibroid and patient-matched myometrial tissues were cut into approximately 5-mm-thick specimens with flat, parallel surfaces. Samples were maintained in serum-free Dulbecco’s Modified Eagle Medium (DMEM) until testing to preserve tissue hydration. Immediately before testing, excess surface medium was gently removed using a Kimwipe, and each specimen was placed on a Criterion Model 42 Universal Testing System (MTS Systems Corporation, Eden Prairie, MN, USA) equipped with a 5-N load cell. The upper compression platen was lowered until full contact with the tissue was achieved under minimal preload, after which samples were allowed to equilibrate for 1 min. Unconfined compression testing was performed at a displacement rate of 2.0 mm min⁻¹ until 10% engineering strain was reached. Young’s modulus was calculated as the slope of the linear region of the stress–strain curve over the 5–10% strain interval.

### Tissue processing and histological staining

Tissues were fixed in 10% neutral buffered formalin (Sigma-Aldrich HT501320) for 48 h, followed by stepwise dehydration in 70%, 80%, 95%, and 100% ethanol followed by xylenes (ThermoFisher Scientific X3P-1GAL) for 1 h and stored in paraffin wax overnight at 55-60 °C. Tissue were sectioned by JHU Oncology Tissue Services SKCCC core facility at a thickness of 4 μm per slide.

Formalin-fixed paraffin-embedded (FFPE) sections of fibroids and matched myometrium from 14 patients were stained for hematoxylin and eosin (H&E) and Masson’s Trichrome by the JHU Oncology Tissue Services SKCCC core facility. Slides were imaged with NanoZoomer-XR digital slide scanner (Hamamatsu Photonics) and viewed using NDP.view2 software (Hamamatsu Photonics, v2.9.25).

### Immunofluorescence staining

FFPE sectioned slides were baked in an oven, deparaffinized in xylene and rehydrated using decreasing concentrations (100%, 95%, 80%, 70%) of ethanol and rinsed with Type I water. Antigen retrieval was performed by adding slides into heated 1x AR6 Buffer (Akoya Biosciences AR600250ML) for 20 min in a steamer. The slides were rinsed with Type 1 water, and endogenous peroxidases quenched using fresh 3% H_2_O_2_ for 15 minutes. Slides were then blocked in 10% BSA in 1x Tris Buffered Saline with Tween20 (TBS-T) (Cell Signaling Technology 9997S) with 0.05% Tween 20 for 30 min and incubated with the target primary antibody listed in **Supplementary Table 34** for 30 min at RT. Sections incubated without primary antibody (primary delete) were used as negative controls to assess nonspecific secondary antibody binding. Aftewards, slides were washed with 1x TBS-T and incubated with corresponding horseradish peroxidase (HRP) polymer-conjugated secondary antibody (Biocare Medical) for 30 min. Then, slides were washed with 1x TBS-T and incubated in Opal 570 Reagent (1:150) (Akoya Biosciences, FP1488001KT) or Opal 650 Reagent (1:150) (Akoya Biosciences, FP1496001KT) in 1x Plus Amplification Diluent (Akoya Biosciences, FP1498) for 10 minutes in the dark. Slides were then washed in 1× TBS-T, rinsed with Type I water, and finally incubated with 1× Spectral DAPI (Akoya Biosciences, FP1490) for 5 min. DAKO Fluorescence Mounting Medium (Agilent, S302380-2) and coverslips were applied and slides were allowed to dry overnight in the dark. Slides were imaged and stitched using the Axio Imager A2 microscope (Carl Zeiss Microscopy, LLC, model #: 490022-0009-000) and Zeiss ZEN 3.4 (blue edition) software.

### Machine learning-based 3D reconstruction

We performed three-dimensional (3D) reconstruction of serial FFPE sections from one patient-derived fibroid and its matched myometrium using CODA [79], a deep learning algorithm for 3D tissue mapping of histological sections. H&E-stained serial FFPE sections were digitized at 20x magnification (pixel size = 0.4543 µm/pixel) using a NanoZoomer-XR digital slide scanner. Serial whole-slide H&E images were first aligned using rigid-body registration, followed by deformation-field image registration to correct for inter-slice deformation.

A deep learning–based semantic segmentation model using the DeepLabv3+ framework [80] was trained on manually annotated H&E images to classify tissue compartments at a downsampling factor of 4 (1.81 µm spatial resolution relative to the scanned H&E images). Annotated classes included smooth muscle, dense ECM, loose ECM, vasculature, and background. The trained model was applied to the serial H&E image stack to generate tissue compartment maps, which were subsequently used to reconstruct the three-dimensional microarchitecture of the fibroid and matched myometrium.

### HALO cell counting

Cell counting was performed using the image analysis platform HALO (version 3.3.2541.285; Indica Labs), using Spatial Analysis module. Cell nuclei (DAPI-positive cells) were defined using nuclear contrast threshold, minimum nuclear intensity, maximum image brightness, nuclear segmentation aggressiveness, and nuclear size. From DAPI-positive cells, p16-positive cells were selected using the nucleus positive threshold. The percentage was calculated as the number of cells with certain phenotype from the total number of DAPI-positive cells.

### Tissue composition and vascular lumen quantification from MTS-stained sections

Masson’s trichrome stained (MTS)-sections from fibroid (n=10) and matched myometrial (n=8) tissues were digitized and analyzed using QuPath (version 0.6.0). Whole tissue sections were analyzed without manual region-of-interest selection. Tissue components were identified using a pixel classification approach (OpenCV Random Trees classifier) trained to distinguish lumen, smooth muscle, and extracellular matrix (ECM) compartments. Feature extraction included multiscale Gaussian and Laplacian filters. Following training, the classifier was applied uniformly across all sections using identical parameters. The area of each compartment (lumen, smooth muscle, and extracellular matrix) was quantified and normalized to total tissue area to determine relative tissue composition. For vascular lumen count, lumen regions identified by pixel classification were converted into discrete objects for object-based quantification. Individual lumen objects were counted without further subclassification. Vascular lumen density was defined as the proportion of total tissue area classified as lumen, and lumen number was defined as the total count of lumen objects per section. Statistical comparisons between fibroid and matched myometrial samples were performed using a paired two-sided t-test.

### p16⁺ cell detection and subregion identification

Cell detection was performed on whole-slide images using StarDist with a 10 µm dilation coefficient, applied to the DAPI channel within QuPath v0.5.1. Following cell detection, cells were classified as p16⁺ based on their staining intensity. Specifically, cells were designated as p16⁺ if their intensity values exceeded the sum of the mean and standard deviation (mean + SD) of all detected cells. To identify spatially distinct subregions within the tissue, we extracted the centroid positions of all detected cells and applied K-means++ clustering, identifying eight subregions. Clustering was performed using the Euclidean distances between cell centroids to group spatially coherent regions.

### Statistics

Statistical methods used for scRNA-seq analysis throughout the study are described in the Methods sections, corresponding figure legends, and Main text. All other statistical analyses were performed using GraphPad Prism (versions 10 and 11). The statistical test used for each analysis and relevant details of data presentation are specified in the corresponding figure legend.

### Data visualization

Schematics used in figures were created with BioRender.com. Volcano plots were created using EnhancedVolcano v1.16.0, and chord diagrams were constructed using circlize v0.4.16 [81] and refined in Affinity Designer. 3D CODA tissue visualizations were generated using 3D Slicer (v5.6.2) [82]. Heatmaps were created using ComplexHeatmap v2.14.0 or ggplot2 from the tidyverse v2.0.0. Network communication plots were produced with igraph. Supplementary packages included ggnewscale v0.4.10, ggrepel v0.9.5, and svglite v2.1.3 for visualization, and stringdist v0.9.12, reshape2 v1.4.4, and plyr v1.8.9 for data manipulation.

## Data availability

All sequencing data generated in this study will be deposited in the NCBI Gene Expression Omnibus (GEO) prior to publication. The GEO accession number(s) will be provided once available.

## Code availability

Code for all analyses will be made publicly accessible on the Elisseeff lab’s Github repository at www.github.com/Elisseeff-Lab/ upon publication.

## Author contributions

JCM, NC, SN, EJF, MAB, JH and JHE conceptualized and designed the studies. JCM, NC, KBS, HHE, ASR, MSI, MAB, ANR, AF, CF, AN, AR and NR developed and performed experimental work and data analysis. SS, CC, HY, CM and KK performed bioinformatics analyses and SN developed software. MAB, RM, MES, SA, MSI, and SES enrolled patients, collected clinical sample. JCM, SN, NC, and JHE interpreted results. EJF supervised software development. MAB, JS, EJF, JMP and JHE supervised the study design and concept. NC, JCM, and JHE wrote the manuscript. All authors contributed to original draft or reviewing and editing the manuscript and approve of the final form.

## Acknowledgements

We would like to thank C M Cherry Consulting for processing the scRNAseq data, the Johns Hopkins University Transcriptomics and Sequencing core facility for performing scRNA-seq, the JHU Oncology Tissue Services SKCCC core facility, supported by the National Cancer Institute Cancer Center Support Grant P30 CA006973, for histology. This work was supported by NIH Pioneer Award DP1AR076959 (JHE), NIH R01HD094380 (MAB), NIH R01HD111243 (JHE, MAB, and JS), NIH U54AG079779 (EJF, JHE), NIH UH3CA275681 (PHW, JMP), NIH K99AG081564 (JCM), the L’Oréal USA For Women in Science Award (JCM), and NIGMS MIRA grant R35GM157099 (JMP). Additional support was provided by the University of Maryland MPower and PIVOT awards (EJF); this research was supported by the University of Maryland, Baltimore through a grant from the Pivot Strategic Investment Initiative The content is solely the responsibility of the authors and does not necessarily represent the official views of the University of Maryland, Baltimore.

## Competing interests

J.H.E. holds equity in Unity Biotechnology and Aegeria Soft Tissue and is a consultant for Tessara. E.J.F. was on the scientific advisory board of Resistance Bio/Viosera Therapeutics, a paid consultant for Mestag Therapeutics and Piper Sandler, is on the scientific advisory board of the V Foundation, and has a familial relationship to the founder of PushCART Therapeutics.

---

**Extended Data Fig. 1.**
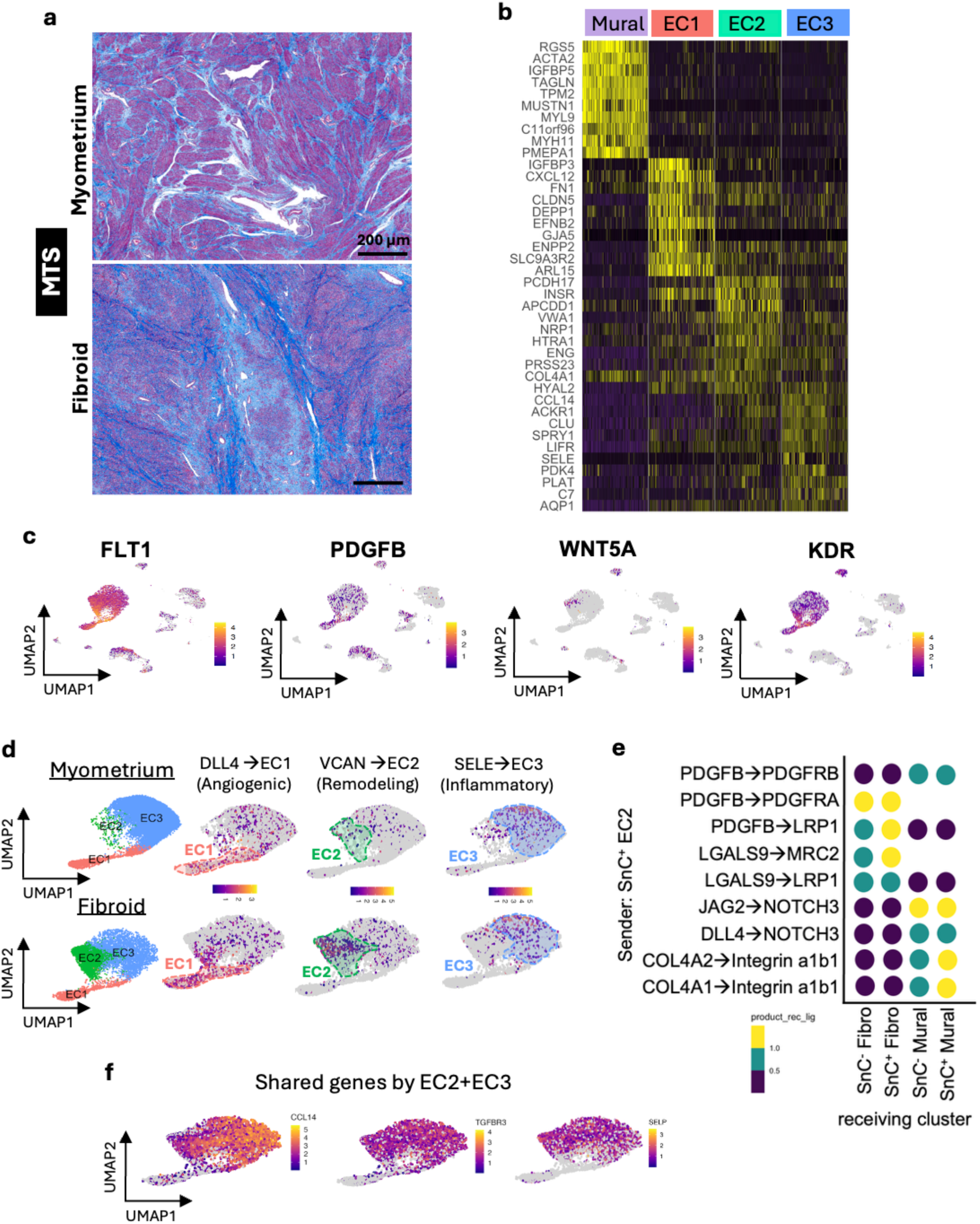
Endothelial heterogeneity and signaling programs in fibroids and matched myometrium. **(a)** Masson’s trichrome staining (MTS) of myometrium and fibroid tissues showing increased extracellular matrix deposition in fibroids. Scale bar, 200 μm. **(b)** Heatmap of marker gene expression defining mural, EC1, EC2, and EC3 populations in fibroids. **(c)** UMAP feature plots showing the expression of representative genes (*FLT1, PDGFB, WNT5A* and *KDR*) across endothelial cell populations in fibroids. **(d)** UMAP projections showing angiogenic (*DLL4*, EC1), remodeling (*VCAN*, EC2) and inflammatory (*SELE*, EC3) programs in myometrium and fibroids. **(e)** Predicted ligand– receptor interactions from SenSig⁺ EC2 cells to SenSig⁺ and SenSig⁻ fibroblast and mural populations, highlighting *PDGF*-, *LGALS9*-, *NOTCH*- and integrin-associated signaling. The interactions shown represent a selected subset of the broader analysis presented in **Supplementary** Fig. 13. **(f)** UMAP feature plots showing genes shared by EC2 and EC3 populations, including *CCL14*, *TGFBR3* and *SELP*.

**Extended Data Fig. 2.**
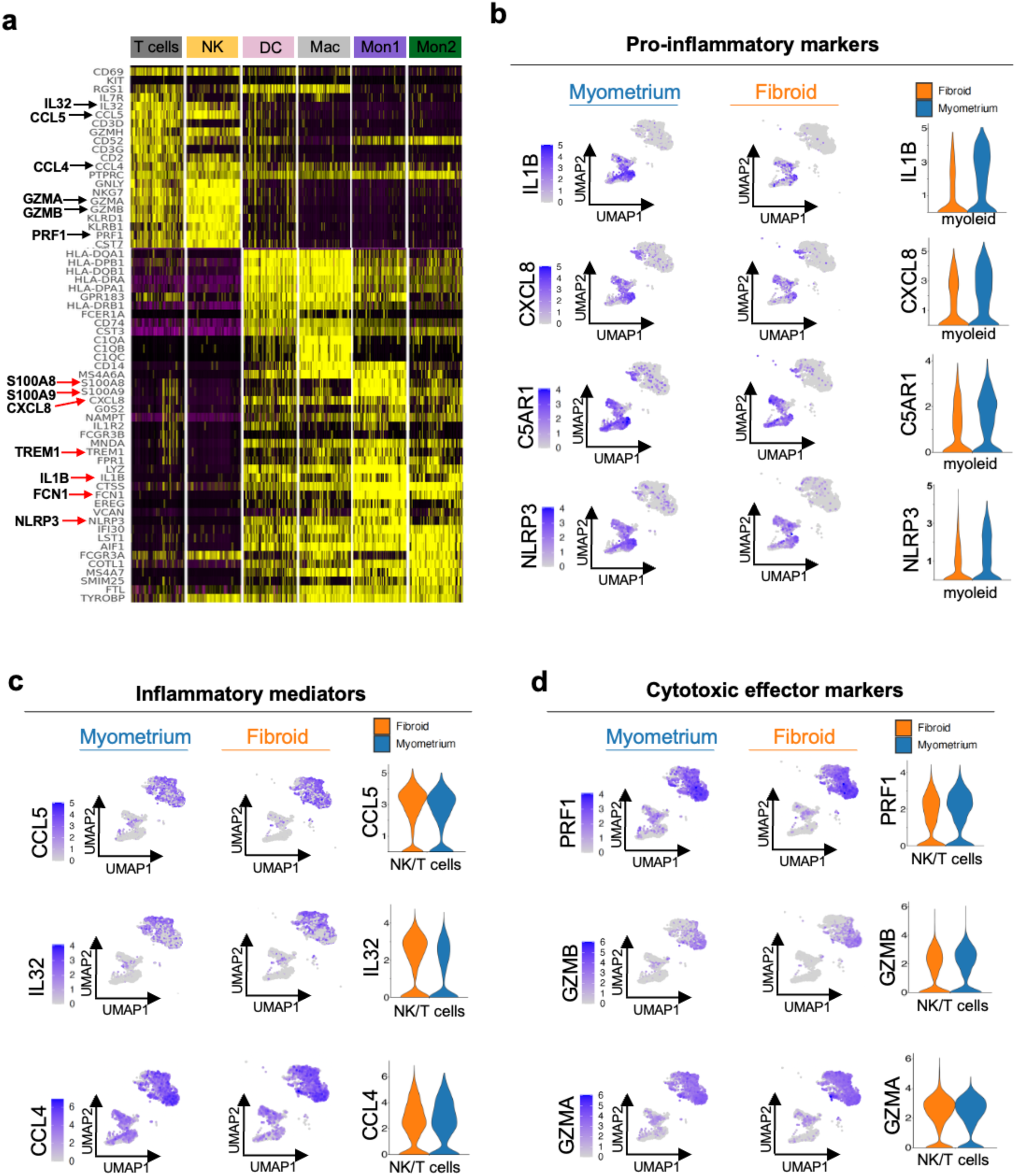
Inflammatory and cytotoxic gene-expression programs in immune cells from human fibroid and myometrium. **(a)** Heatmap showing the expression of selected immune-lineage and activation-associated genes across T cells, natural killer (NK) cells, dendritic cells (DC), macrophages (Mac), and two monocyte subsets (Mon1 and Mon2). Black and red arrows highlight representative lymphoid-associated and myeloid inflammatory genes, respectively. **(b)** UMAP feature plots showing *IL1B*, *CXCL8*, *C5AR1*, and *NLRP3* expression in immune cells from myometrium and fibroid tissues. Corresponding violin plots compare expression distributions within the myeloid compartment. **(c)** UMAP feature plots showing expression of the inflammatory mediators *CCL5*, *IL32*, and *CCL4* in immune cells from myometrium and fibroid tissues, together with corresponding violin plots for the NK/T-cell compartment. **(d)** UMAP feature plots and corresponding violin plots showing expression of the cytotoxic effector genes *PRF1*, *GZMB*, and *GZMA* in the NK/T-cell compartment. Violin plots show fibroid in orange and myometrium in blue.

