## Supplementary Figures for "Senescent cell networks link matrix remodeling and vascular dysfunction in human fibroids"

### Supplementary Figure 1

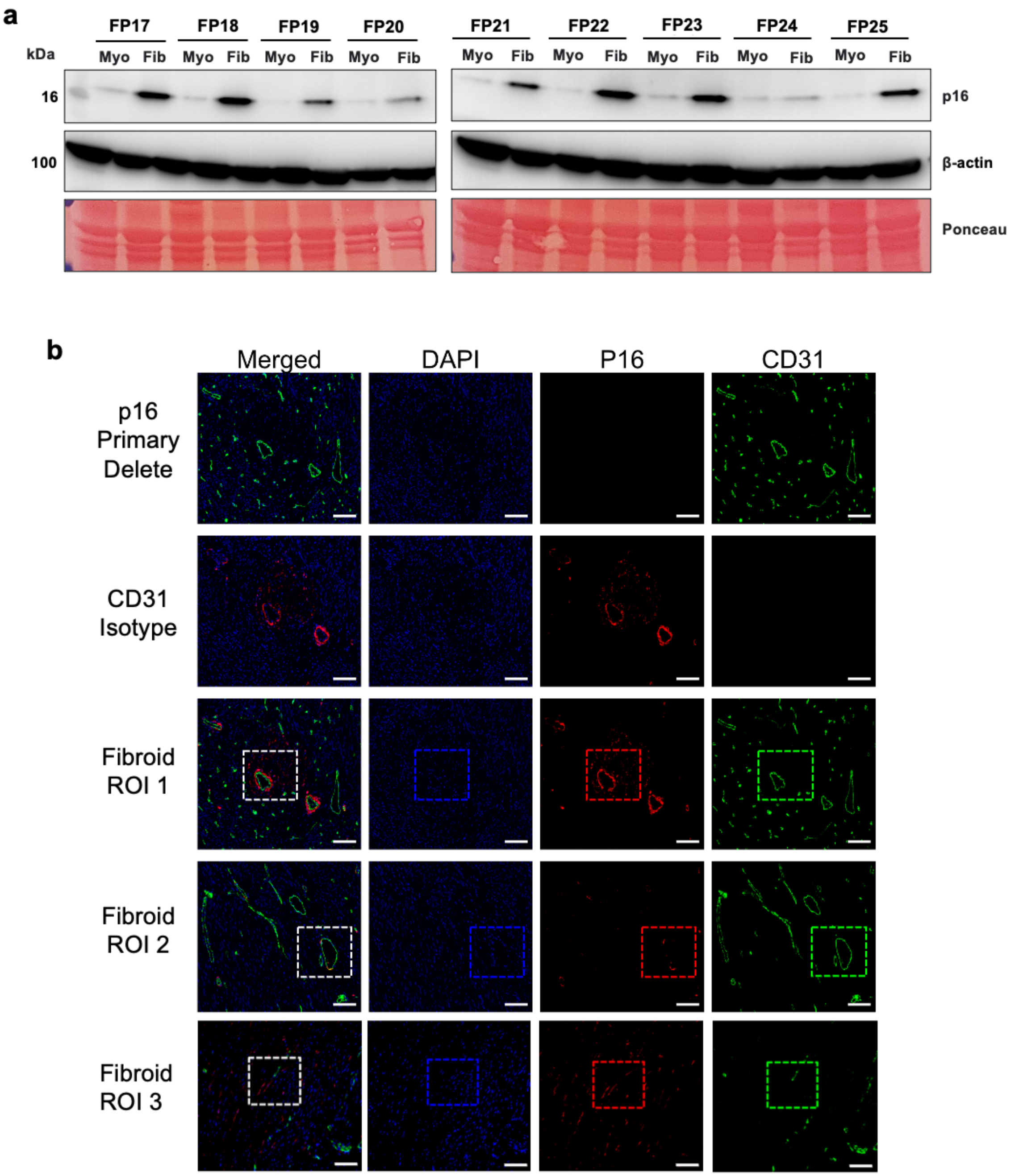


**Supplementary Fig. 1. p16 protein expression in uterine fibroids. (a)** Western blot analysis of p16 protein expression in paired myometrium (Myo) and fibroid (Fib) tissues from patients FP17–FP25. β-actin was used as the control, and Ponceau staining  shows total protein loading. **(b)** p16 SnC cell heterogeneity observed across IF of DAPI (nuclei, blue), p16 (senescence, red), and CD31 (vessels, green) in fibroid samples. Scale bar, 100 µm. The boxed region highlights the area shown in higher magnification in Fig. 1c. Images were linearly contrasted for clarity.

**Supplementary Figure 2**

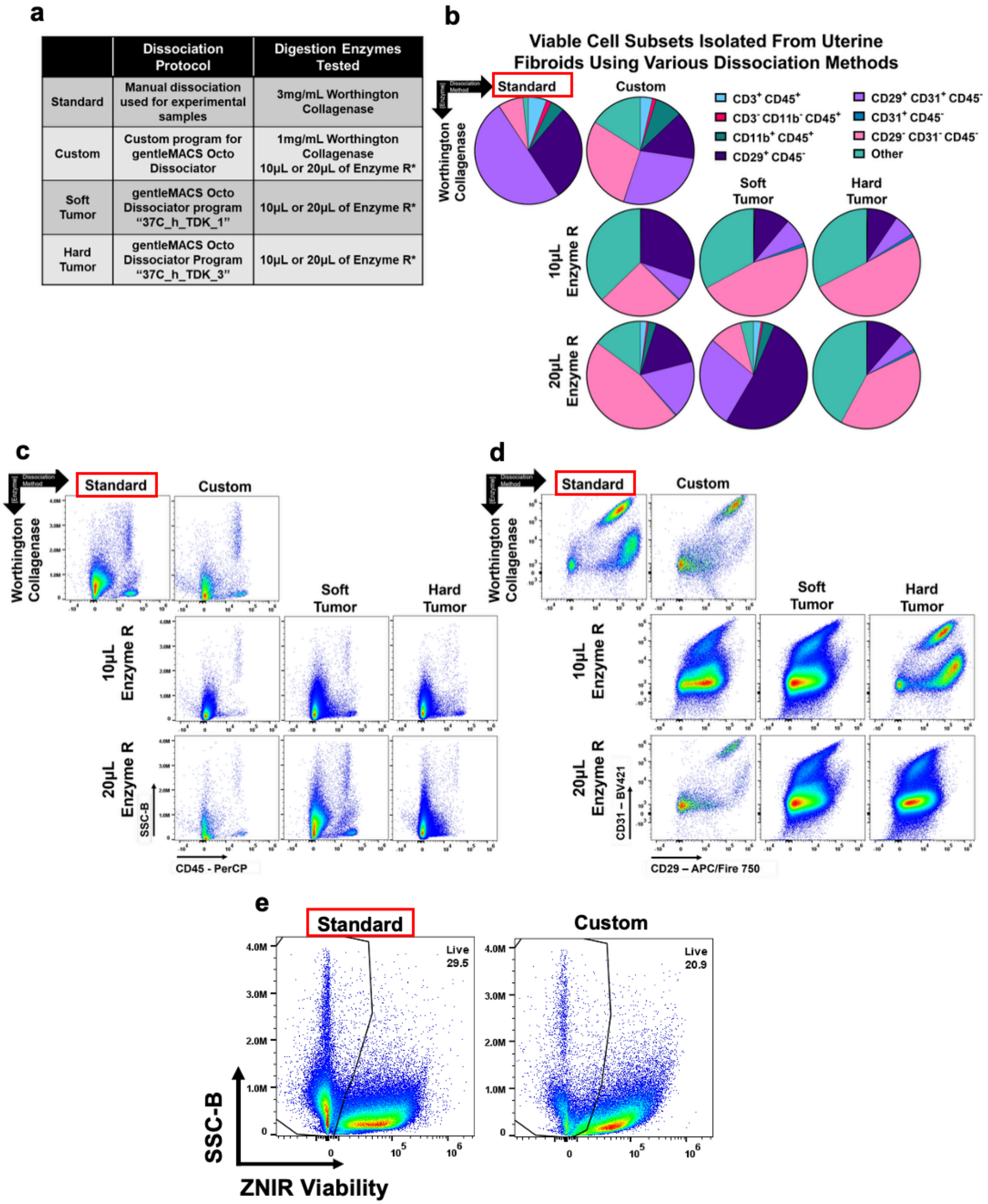

**Supplementary Fig. 2. Optimization of tissue dissociation conditions for recovery of stromal and immune cell populations from human uterine fibroids. (a)** Schematic overview of the four tissue dissociation protocols evaluated for isolation of viable cells from human uterine fibroids, including the standard manual protocol and three gentleMACS Octo Dissociator-based protocols using different enzymatic digestion conditions. **(b)** Relative frequencies of viable immune and stromal cell populations recovered following each tissue dissociation protocol, as determined by flow cytometry. **(c)** Representative flow cytometry pseudocolor plots showing $CD45^{+}$ following each tissue dissociation protocol. Populations were pre-gated to exclude debris, doublets, and non-viable cells. **(d)** Representative flow cytometry pseudocolor plots showing $CD29^{+}$ and $CD31^{+}$ expression within the $CD45^{-}$ stromal compartment following each tissue dissociation protocol. Populations were pre-gated to exclude debris, doublets, and non-viable cells. **(e)** Comparison of cell viability following the standard and custom tissue dissociation protocols. Together, these results demonstrated that the standard manual dissociation protocol preserved recovery of diverse stromal and immune cell populations while providing higher cell viability and was therefore selected for all subsequent flow cytometry and scRNA-seq experiments.

**Supplementary Figure 3**

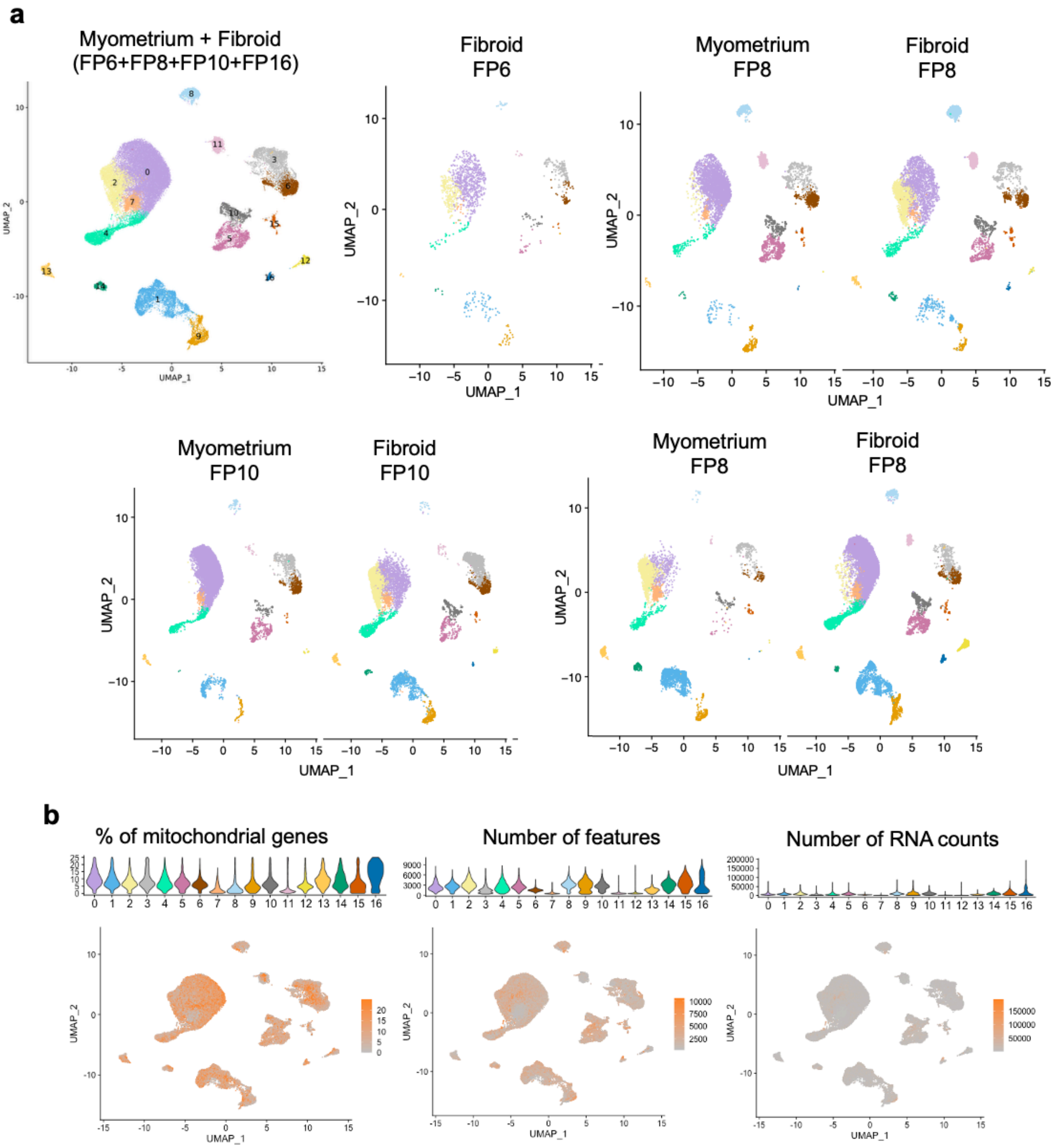


**Supplementary Fig. 3. scRNA-seq profiling of human uterine fibroids and matched myometrium. (a)** UMAP visualization of each patient sample with cell clusters. **(b)** Violin plots illustrating the quality control metrics used for scRNA-seq, including the percentage of mitochondrial gene expression, the number of detected features, and the number of RNA counts per cell across clusters. Corresponding UMAPs show the distribution of each quality control metric.

**Supplementary Figure 4**

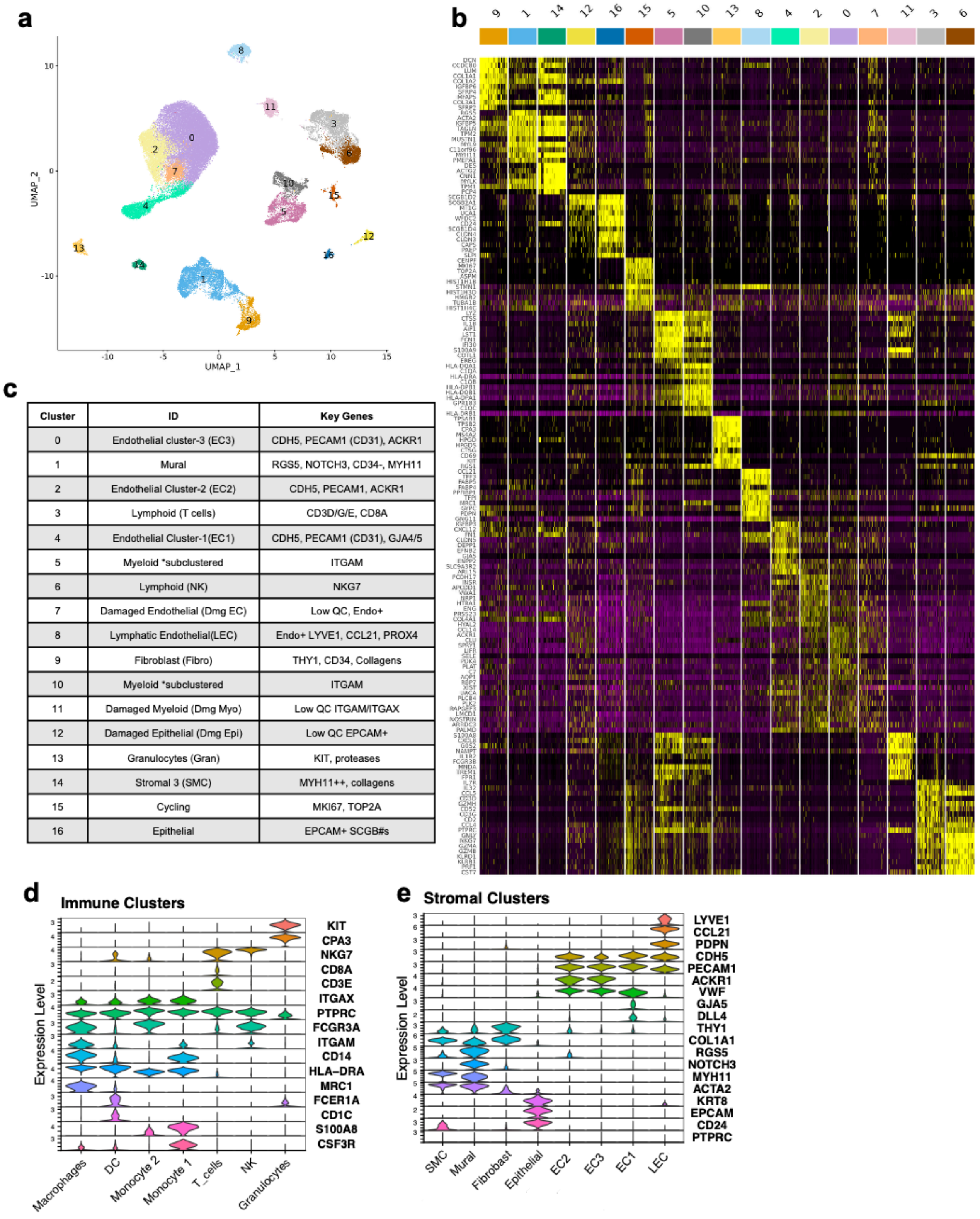


| Cluster | ID | Key Genes |
|---|---|---|
| 0 | Endothelial cluster-3 (EC3) | CDH5, PECAM1 (CD31), ACKR1 |
| 1 | Mural | RGS5, NOTCH3, CD34-, MYH11 |
| 2 | Endothelial Cluster-2 (EC2) | CDH5, PECAM1, ACKR1 |
| 3 | Lymphoid (T cells) | CD3D/G/E, CD8A |
| 4 | Endothelial Cluster-1(EC1) | CDH5, PECAM1 (CD31), GJA4/5 |
| 5 | Myeloid *subclustered | ITGAM |
| 6 | Lymphoid (NK) | NKG7 |
| 7 | Damaged Endothelial (Dmg EC) | Low QC, Endo+ |
| 8 | Lymphatic Endothelial(LEC) | Endo+ LYVE1, CCL21, PROX4 |
| 9 | Fibroblast (Fibro) | THY1, CD34, Collagens |
| 10 | Myeloid *subclustered | ITGAM |
| 11 | Damaged Myeloid (Dmg Myo) | Low QC ITGAM/ITGAX |
| 12 | Damaged Epithelial (Dmg Epi) | Low QC EPCAM+ |
| 13 | Granulocytes (Gran) | KIT, proteases |
| 14 | Stromal 3 (SMC) | MYH11++, collagens |
| 15 | Cycling | MKI67, TOP2A |
| 16 | Epithelial | EPCAM+ SCGB#s |

**Supplementary Fig. 4. scRNA-seq profiling of marker genes for the human fibroid and myometrium. (a)** UMAP visualization showing integrated clustering of high-quality cells from human fibroids (*n* = 4) and matched myometrium (*n* = 3). **(b)** Heatmap indicating the relative expression of each identified cell type's top ten differentially expressed genes. **(c)** List of canonical genes to identify cell clusters. Violin plots showing the expression of canonical marker genes for main cell types for **(d)** immune and **(e)** stromal cell clusters.

**Supplementary Figure 5**

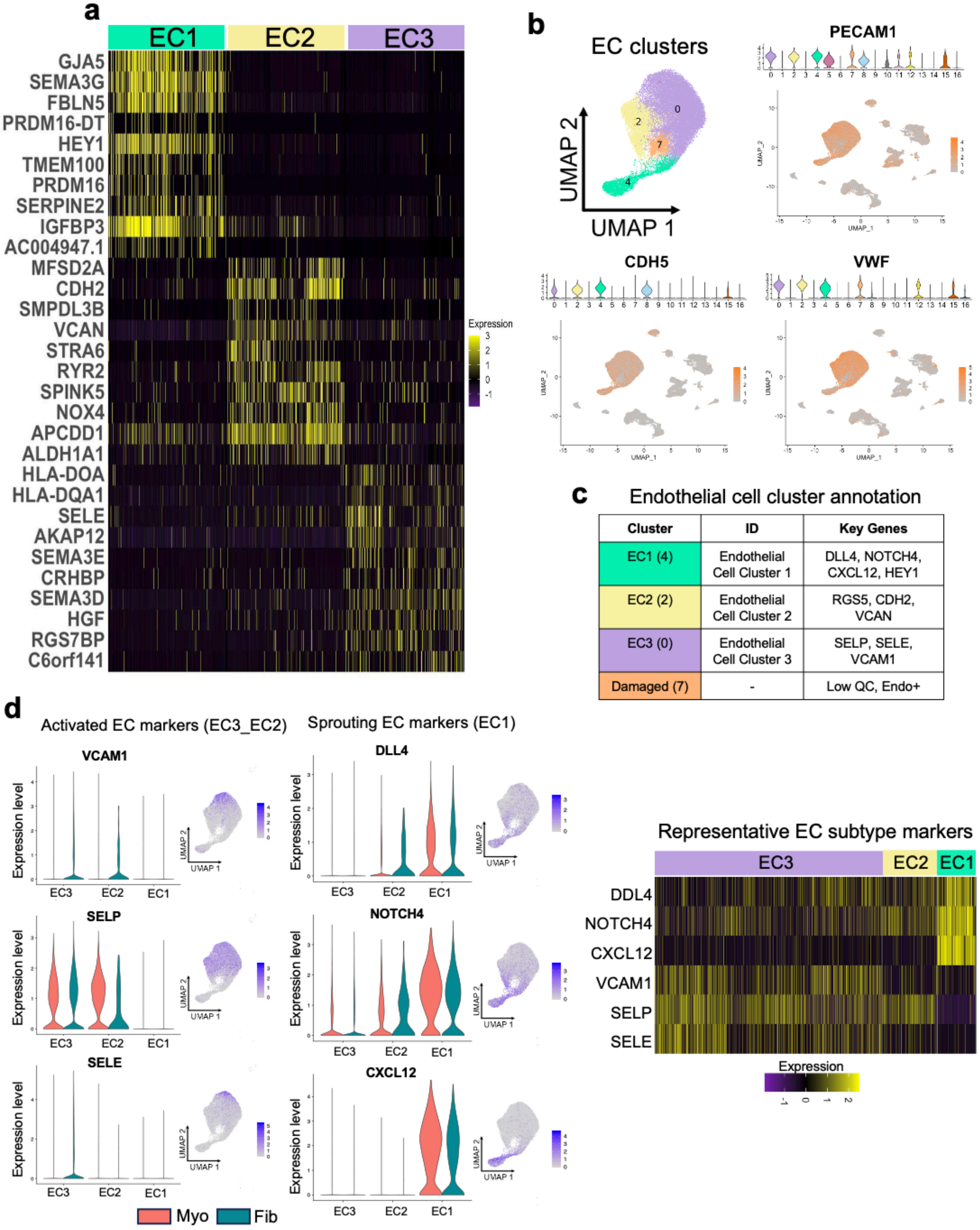


**Supplementary Fig. 5. Clustering of endothelial cells (ECs) from human fibroid and matched myometrium tissues. (a)** Heatmap showing the top 10 differentially expressed genes defining the three endothelial cell subclusters (EC1–EC3) identified from integrated fibroid and matched myometrium samples. **(b)** UMAP visualization of endothelial cells isolated from human myometrium (*n* = 3) and fibroid ($n$ = 4) tissues, identifying three transcriptionally distinct endothelial subclusters (EC1–EC3). Feature plots illustrate the expression of canonical endothelial markers *PECAM1 (CD31)*, *CDH5*, and *VWF*. **(c)** Annotation of endothelial clusters based on their characteristic marker genes and inferred identities. **(d)** Violin plots and feature plots showing representative markers distinguishing endothelial subpopulations. EC1 is enriched for angiogenic/sprouting endothelial markers (*DLL4*, *NOTCH4*, and *CXCL12*), whereas EC2 and EC3 display an activated endothelial phenotype characterized by *VCAM1*, *SELP*, and *SELE* expression. The accompanying heatmap highlights the differential expression of representative EC1, EC2 and EC3 marker genes across endothelial subclusters. Violin plots compare gene expression between myometrium (Myo) and fibroid (Fib).

**Supplementary Figure 6**

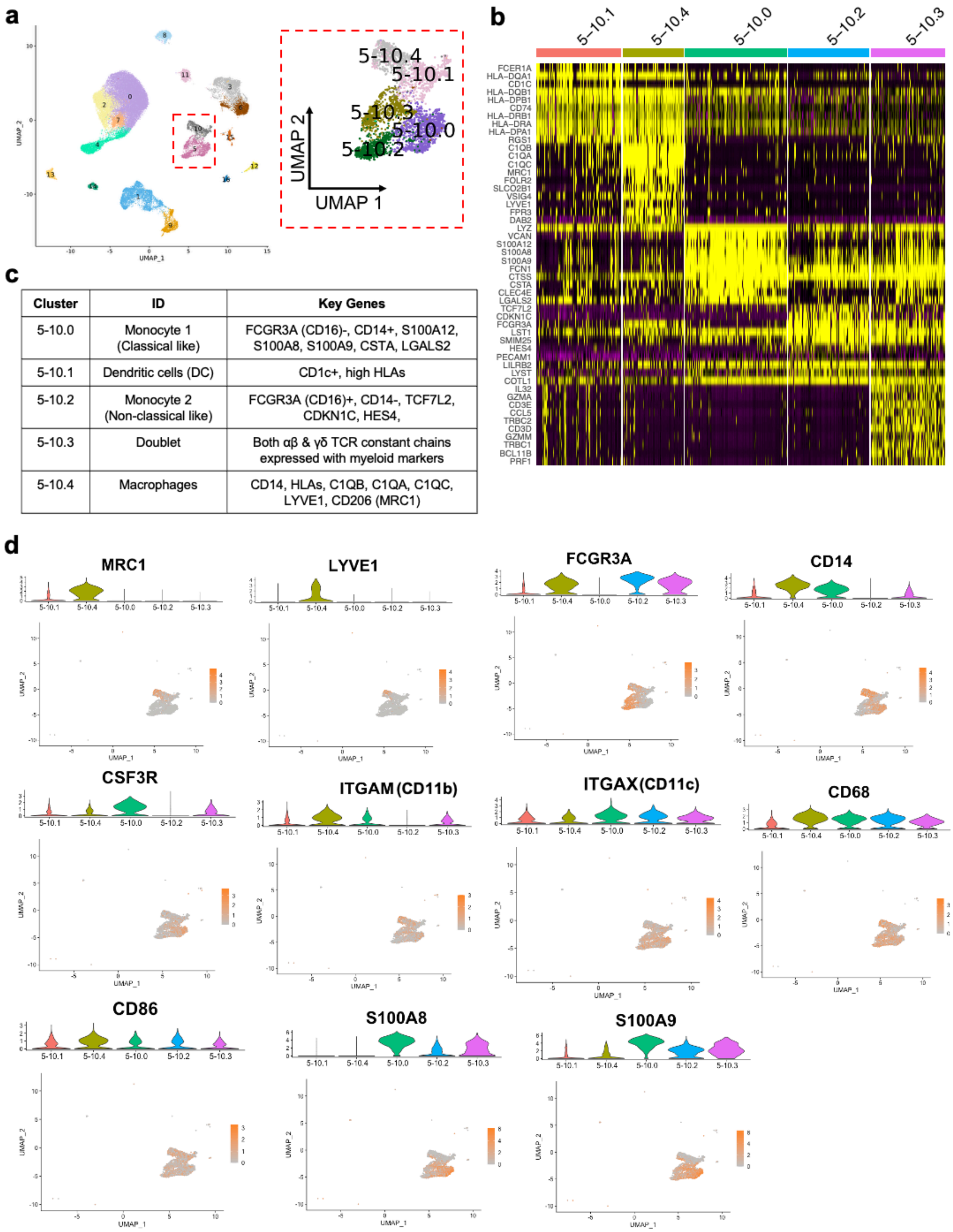


| Cluster | ID | Key Genes |
|---|---|---|
| 5-10.0 | Monocyte 1 (Classical like) | FCGR3A (CD16)-, CD14+, S100A12, S100A8, S100A9, CSTA, LGALS2 |
| 5-10.1 | Dendritic cells (DC) | CD1c+, high HLAs |
| 5-10.2 | Monocyte 2 (Non-classical like) | FCGR3A (CD16)+, CD14-, TCF7L2, CDKN1C, HES4, |
| 5-10.3 | Doublet | Both αβ & γδ TCR constant chains expressed with myeloid markers |
| 5-10.4 | Macrophages | CD14, HLAs, C1QB, C1QA, C1QC, LYVE1, CD206 (MRC1) |

**Supplementary Fig. 6. Subclustering of myeloid cells. (a)** UMAP visualization showing subclustering of myeloid cells from human fibroid (*n* = 4) and matched myometrium (*n* = 3) samples. The inset shows the five identified myeloid subclusters. **(b)** Heatmap showing the top 10 DEGs defining each myeloid subcluster. **(c)** Annotation of myeloid subclusters based on representative marker genes. **(d)** Violin plots and corresponding feature plots showing the expression of representative marker genes used to distinguish macrophages, dendritic cells, classical monocytes, and non-classical monocytes.

**Supplementary Figure 7**

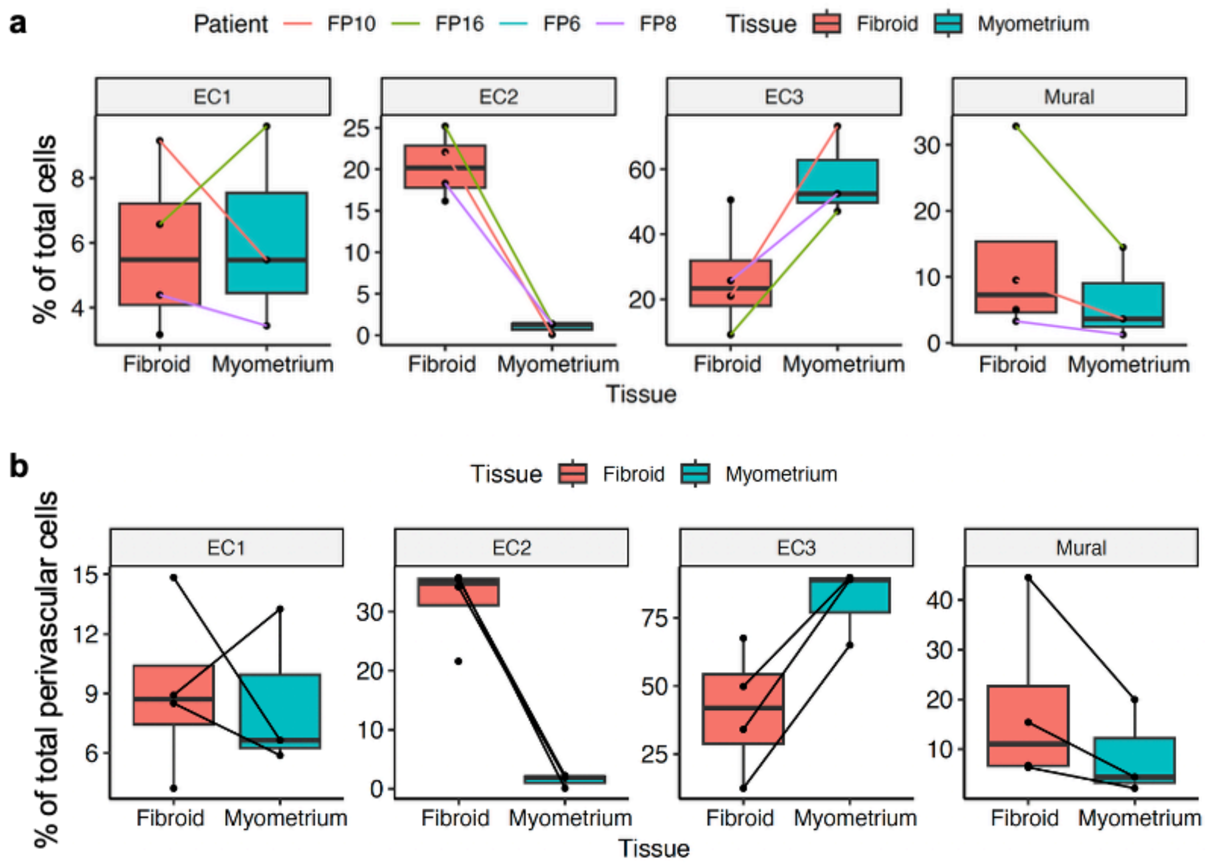


**Supplementary Fig. 7. EC2 endothelial cells are enriched in human uterine fibroids compared with matched myometrium. (a)** Relative abundance of EC1, EC2, EC3, and mural cell populations expressed as a percentage of the total cells in fibroid (*n* = 4) and matched myometrium (*n* = 3) samples. Boxplots show the distribution across samples, points represent individual patient samples, and lines connect the three matched fibroid–myometrium pairs. Differential abundance between fibroid and matched myometrium samples was assessed using an edgeR quasi-likelihood F-test with patient identity included in the model, followed by false discovery rate (FDR) correction. EC2 was significantly enriched in fibroids (FDR = 0.0004). **(b)** Relative abundance of EC1, EC2, EC3, and mural cell populations expressed as a percentage of the total perivascular cell population in fibroid (*n* = 4) and matched myometrium (*n* = 3) samples. Boxplots show the distribution across samples, points represent individual patient samples, and lines connect the three matched fibroid–myometrium pairs.

**Supplementary Figure 8**

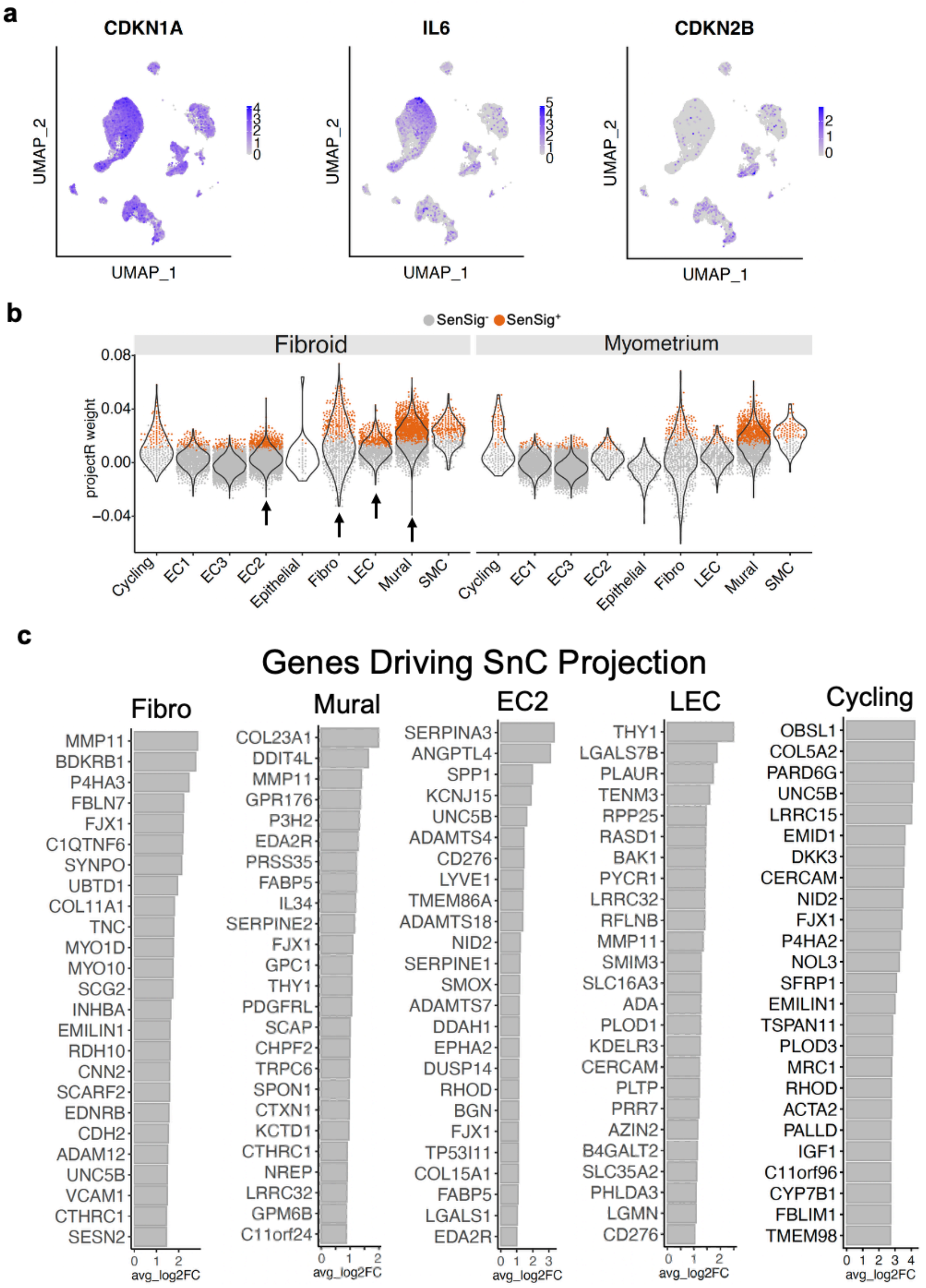


**Supplementary Fig. 8. Senescence-associated transcriptional programs across fibroid and myometrium single-cell datasets.** (a) Feature plots showing expression of canonical senescence markers, including CDKN2B (p15), CDKN1A (p21), and SASP-associated genes such as IL6, across all cell populations. These markers exhibit limited and non-specific expression patterns. (b) Violin plot showing projection of the SenSig into integrated fibroid and myometrium single-cell transcriptomes using the projectR. Cells with a positive projection weight and p-value less than 0.01 were labeled as SenSig⁺. SenSig⁺ cells are highlighted in orange, with higher projection weights within mural cells, fibroblasts, lymphatic endothelial cells (LEC), and a subset of endothelial cells (EC2 cluster). (c) Genes driving SenSig projection in mural cells, fibroblasts, LEC, and EC2 cluster.

**Supplementary Figure 9**

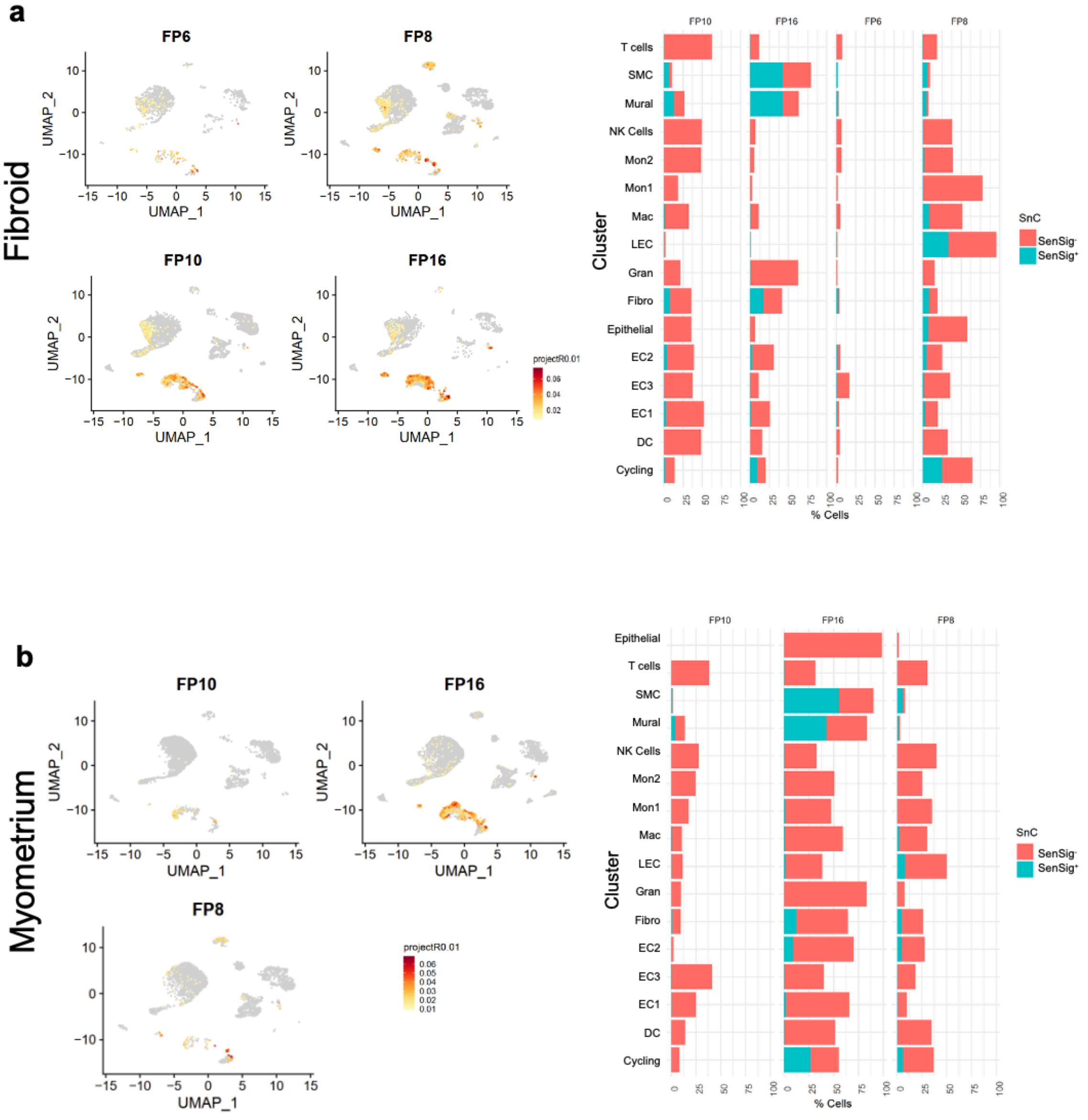


**Supplementary Fig. 9. Patient-specific distribution of SenSig+ cells across cell types in fibroids and matched myometrium.** (**a**) UMAP projections showing the distribution of SenSig+ cells across individual fibroid samples (*n* = 4; FP6, FP8, FP10 and FP16). (**b**) UMAP projections showing the distribution of SenSig⁺ cells across matched myometrium samples (*n* = 3; FP8, FP10 and FP16). SenSig⁺ cells were distributed across multiple cell populations, with enrichment in specific clusters in a patient-dependent manner. Bar plots summarize the proportions of SenSig⁺ and SenSig⁻ cells across annotated cell types in each sample.

**Supplementary Figure 10**

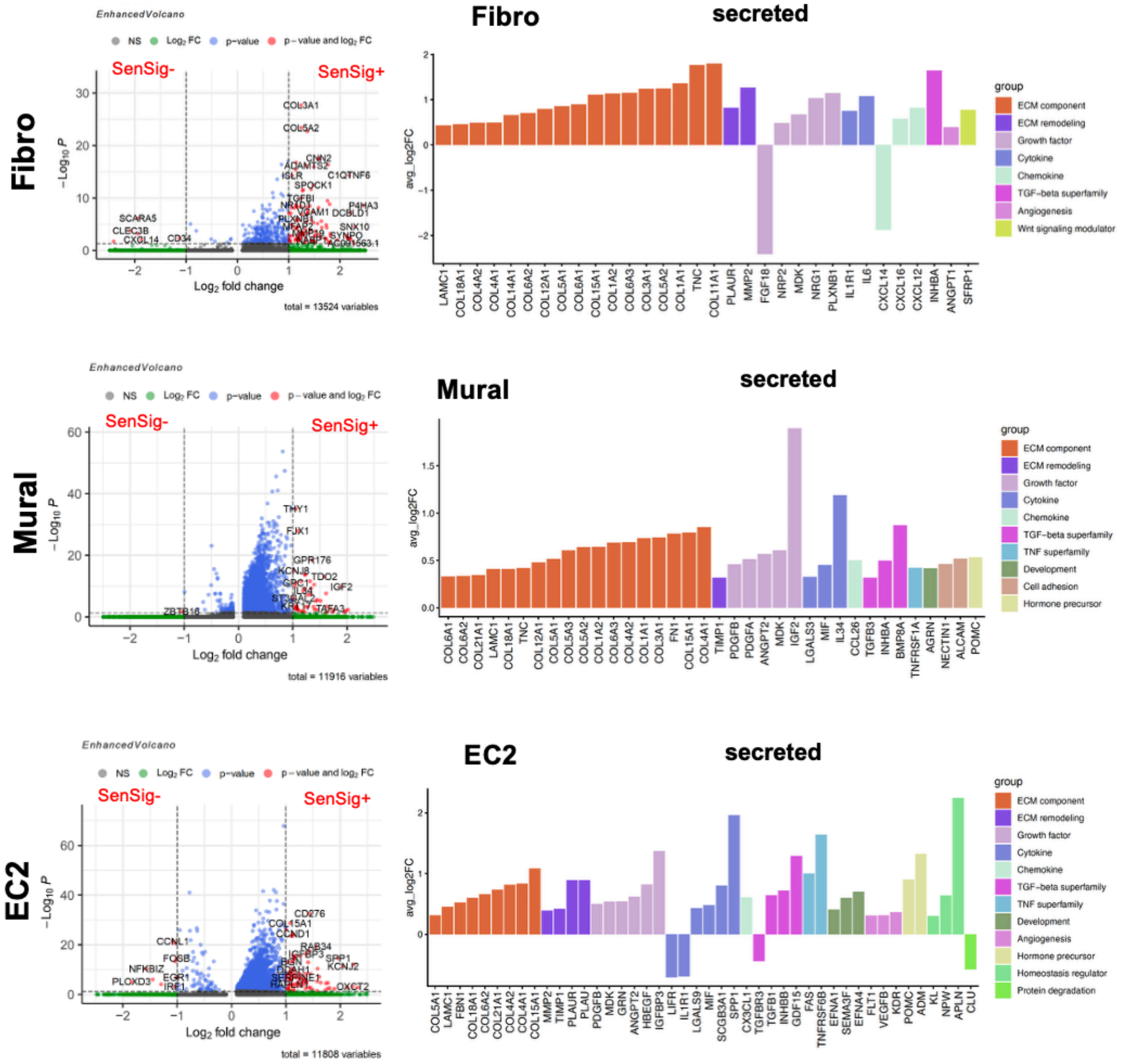


**Supplementary Fig. 10. Differential gene expression and secreted-factor programs in SenSig⁺ and SenSig⁻ stromal populations.** Volcano plots show differential gene expression between SenSig⁺ and SenSig⁻ cells within fibroblast, mural and EC2 populations. Positive $log_2$ fold-change values indicate genes enriched in SenSig⁺ cells, whereas negative values indicate genes enriched in SenSig⁻ cells. Bar plots show selected differentially expressed genes encoding secreted factors (adjusted $P$ value $< 0.05$ and absolute average $log_2$ fold change $> 0.3$), grouped according to the functional categories indicated by color, including extracellular matrix (ECM) components, ECM-remodeling factors, growth factors, cytokines, chemokines, TGF-β superfamily members, angiogenesis-related factors and other functional classes.

**Supplementary Figure 11**

a

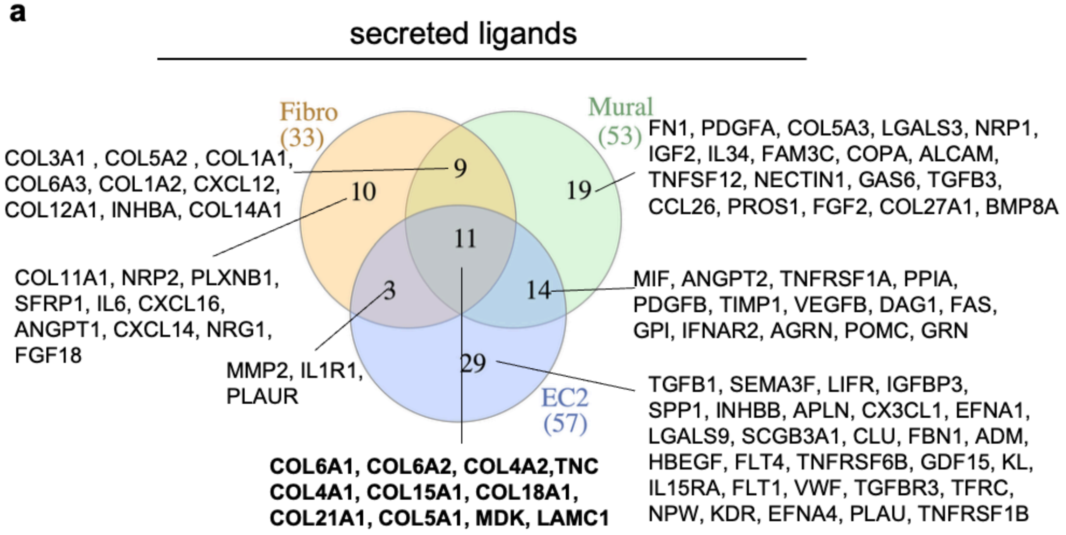


b

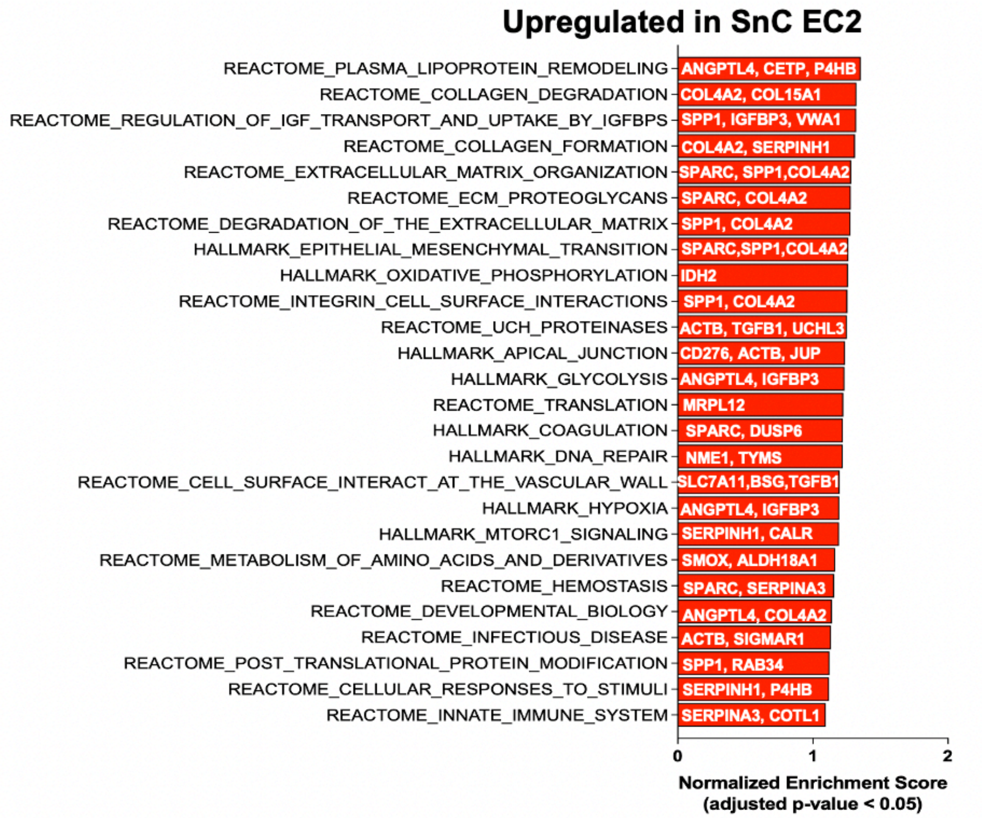


**Supplementary Fig. 11. Shared SASP genes across SnC subtypes and functional pathway enrichment in EC2 cells**. **(a)** Venn diagram showing the numbers of SASP genes shared among and unique to fibroblast, mural and EC2 SnC subtypes. Numbers in parentheses indicate the total number of SASP genes in each subtype, whereas numbers within the circles indicate subtype-specific and shared genes. **(b)** Gene set enrichment analysis (GSEA) comparing SenSig$^{+}$ and SenSig$^{-}$ EC2 cells and showing significantly enriched Hallmark and Reactome pathways (adjusted $P < 0.05$). Bars represent normalized enrichment scores, and labels within the bars indicate selected leading-edge genes.

**Supplementary Figure 12**

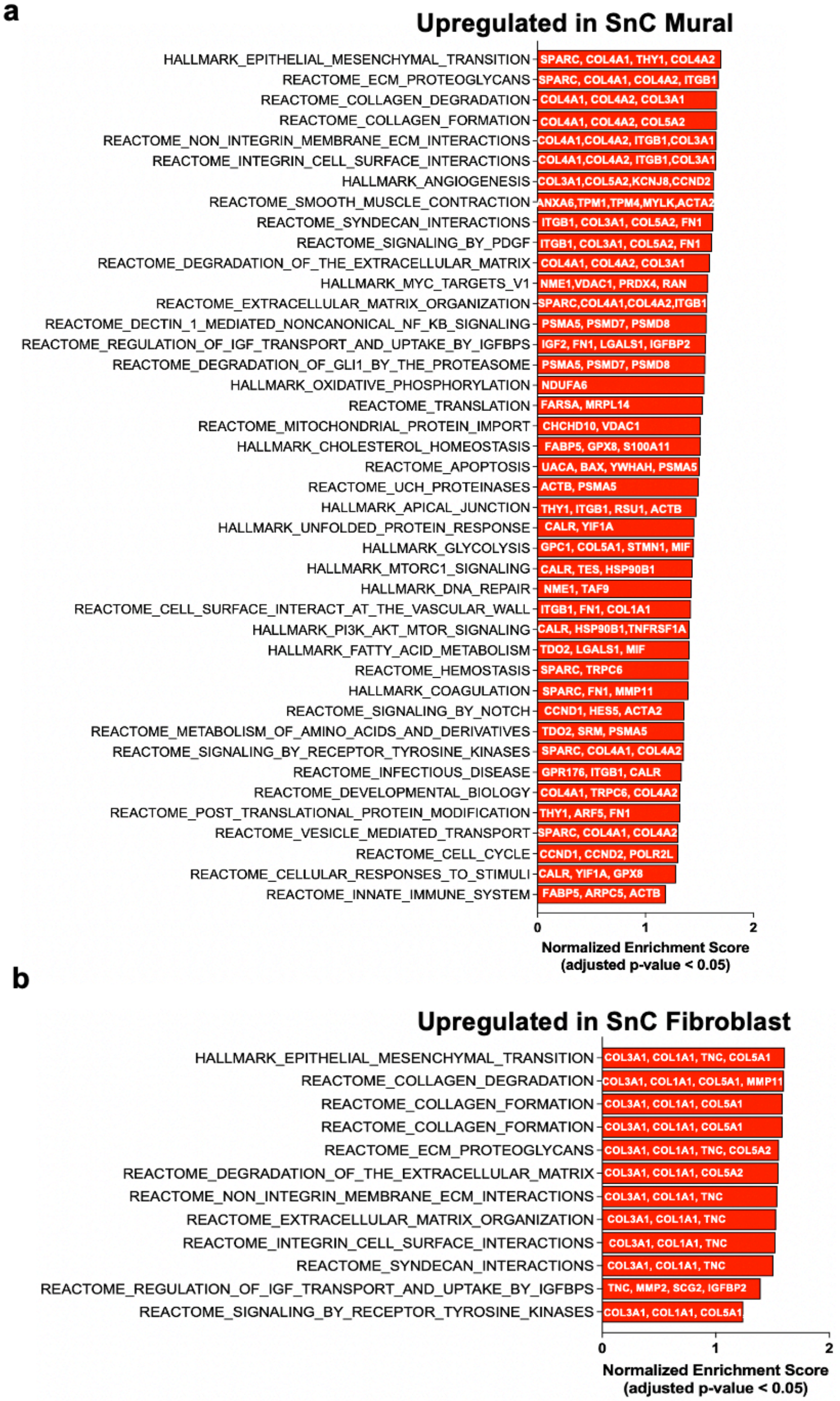


**Supplementary Fig. 12. Functional pathway enrichment in SnC mural cells and fibroblasts. (a)** Gene set enrichment analysis (GSEA) comparing SenSig$^{+}$ and SenSig$^{-}$ mural cells and showing significantly enriched Hallmark and Reactome pathways. **(b)** GSEA comparing SenSig$^{+}$ and SenSig$^{-}$ fibroblasts and showing significantly enriched Hallmark and Reactome pathways. Bars represent normalized enrichment scores (NES), and labels within the bars indicate selected leading-edge genes. Significantly enriched pathways were defined by an adjusted *P* value < 0.05.

**Supplementary Figure 13**

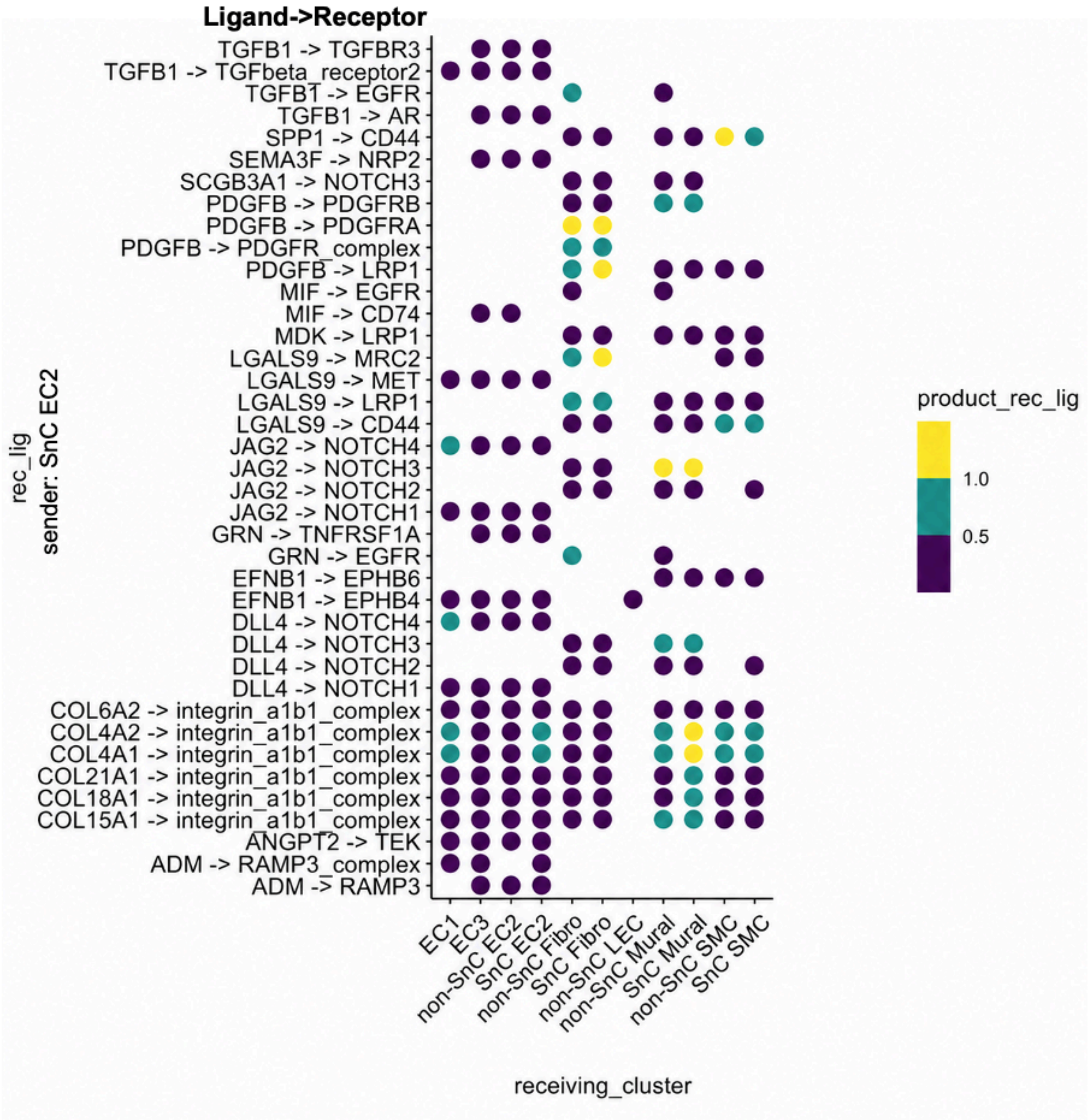


**Supplementary Fig. 13. Predicted ligand–receptor signaling from SnC EC2 cells to vascular and stromal populations in fibroids.** Dot plot showing predicted interactions from SnC EC2 cells to the indicated receiving populations. Candidate ligands were differentially expressed between SnC and non-SnC EC2 cells (adjusted $P < 0.05$; absolute average $\log_2$ fold change $> 0.3$), and interactions detected in at least three of four fibroid samples were retained. Rows indicate ligand–receptor pairs, columns indicate receiving populations and dot color represents the interaction score (product_rec_lig).

**Supplementary Figure 14**

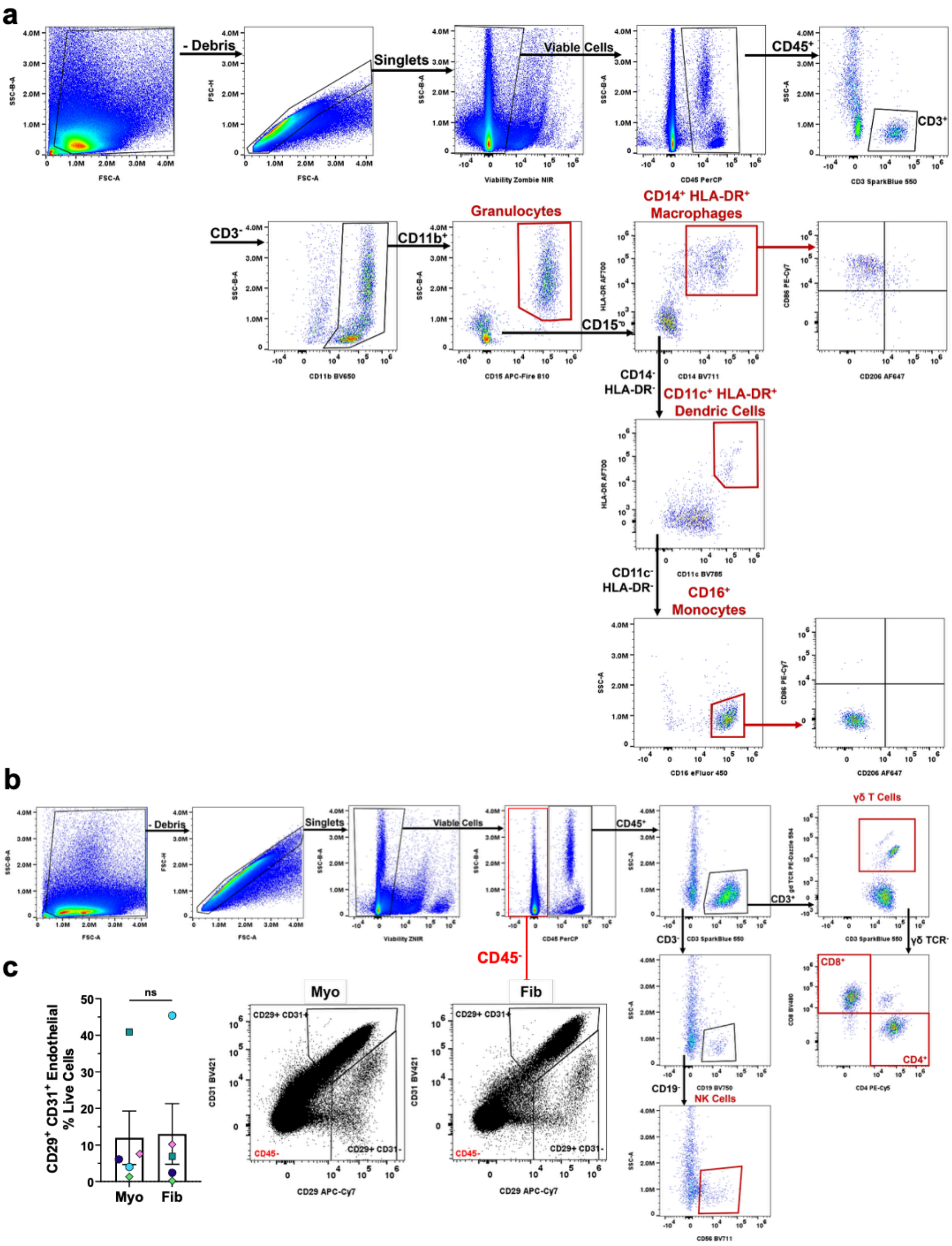


**Supplementary Fig. 14. Gating strategies for flow cytometric analysis and population quantification. (a)** Representative gating strategy for identifying myeloid populations in uterine fibroids. **(b)** Representative gating strategy for identifying lymphoid populations in uterine fibroids. The same gating strategies were applied to available myometrium samples. **(c)** Representative gating and quantification of $CD29^{+}CD31^{+}$ endothelial cells within the viable $CD45^{-}$ population in patient-matched myometrium and fibroid samples. Endothelial-cell frequencies were determined from the CD45⁻ fraction of samples stained with the myeloid flow-cytometry panel. ns, not significant.

**Supplementary Figure 15**

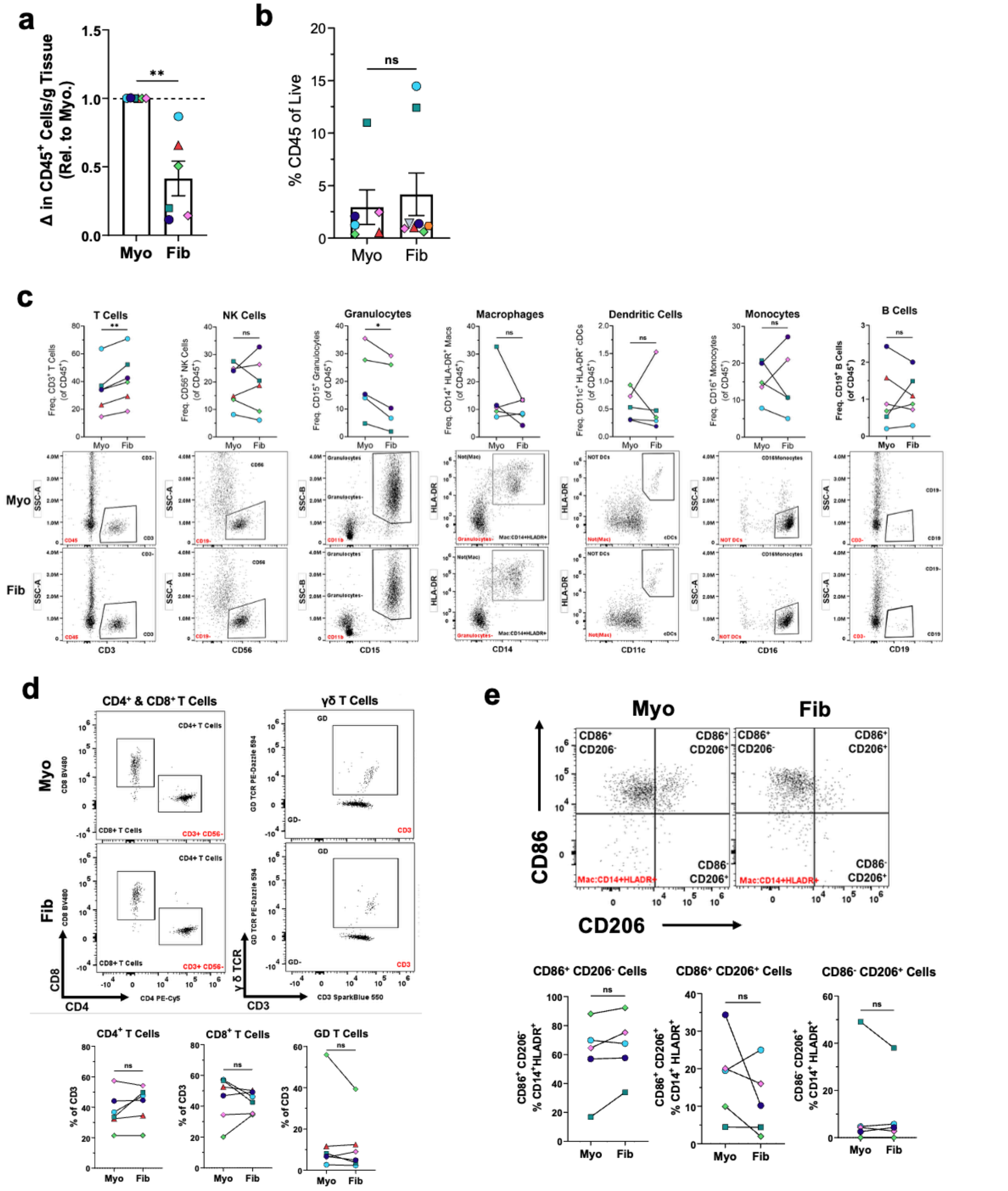

**Supplementary Fig. 15. Flow-cytometric profiling of immune-cell populations in myometrium and fibroid tissues. (a)** Quantification of live $CD45^+$ cells per gram of tissue in fibroids relative to patient-matched myometrium. **(b)** Frequency of $CD45^+$ cells among viable cells in myometrium and fibroid samples. **(c)** Frequencies of the indicated $CD45^+$ immune-cell subsets in patient-matched myometrium and fibroid tissues, with representative flow-cytometry plots shown below. These populations were initially identified by scRNA-seq and subsequently evaluated by flow cytometry. **(d)** Representative plots and quantification of $CD4^+$, $CD8^+$ and γδ T-cell subsets within the $CD3^+$ T-cell population. **(e)** Macrophage phenotypes assessed by CD86 and CD206 expression within $CD14^+HLA\text{-}DR^+$ cells, with quantification of $CD86^+CD206^-$, $CD86^+CD206^+$ and $CD86^-CD206^+$ populations. Myometrium plots were normalized to display the same number of events as the corresponding fibroid plots. Symbols represent individual samples, and lines connect patient-matched myometrium and fibroid samples. Statistical significance was determined using paired tests. ns, not significant; $*P < 0.05$; $**P < 0.01$.

### Supplementary Figure 16

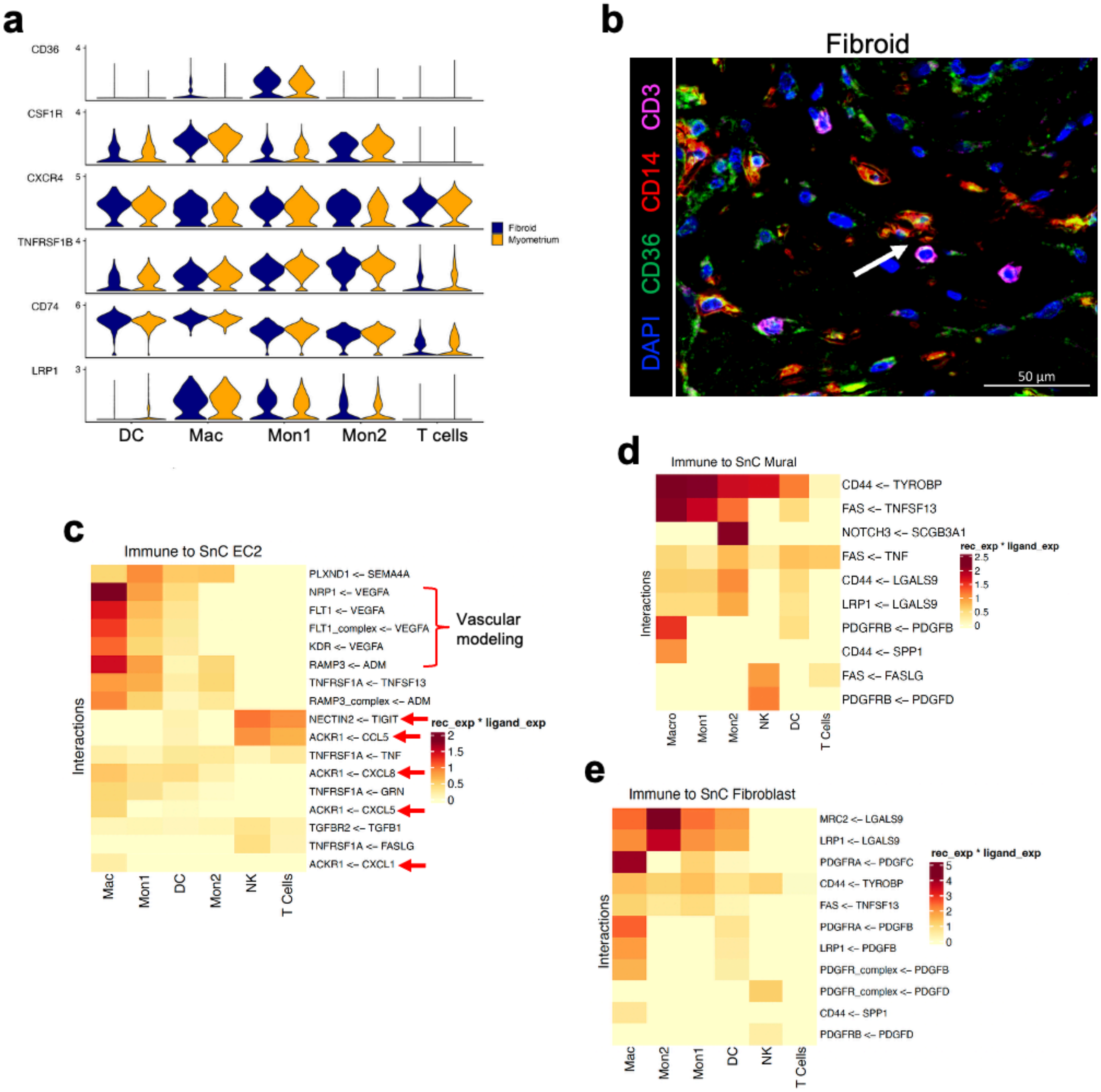


**Supplementary Fig. 16. Immune receptor expression, spatial organization and predicted immune-to-SnC signaling. (a)** Violin plots showing the expression of selected receptors (*CD36*, *CSF1R*, *CXCR4*, *TNFRSF1B*, *CD74* and *LRP1*) across immune populations in fibroid ($n$ = 4) and matched myometrium ($n$ = 3) scRNA-seq datasets. **(b)** Representative IF image of fibroid tissue stained for CD14 (red), CD36 (green), CD3 (magenta) and DAPI (blue). The arrow indicates a CD3$^{+}$ T cell in close proximity to CD14$^{+}$CD36$^{+}$ myeloid cells. Scale bar, 50 µm. **(c–e)** Heatmaps showing predicted ligand–receptor interactions from immune populations to **(c)** SnC EC2, **(d)** SnC mural and **(e)** SnC fibroblast populations in fibroids. Columns indicate immune sender populations, rows indicate receptor–ligand pairs and color represents the product of receptor and ligand expression (rec_exp × ligand_exp). Selected interactions include *VEGFA–FLT1*/*KDR*/*NRP1* and *ADM–RAMP3* signaling to SnC EC2 cells, *LGALS9–LRP1*/*MRC2* signaling and *PDGF* ligand–receptor interactions with SnC mural and fibroblast populations. Mac, macrophages; Mon1 and Mon2, monocyte subsets; DC, dendritic cells; NK, natural killer cells.

**Supplementary Figure 17**

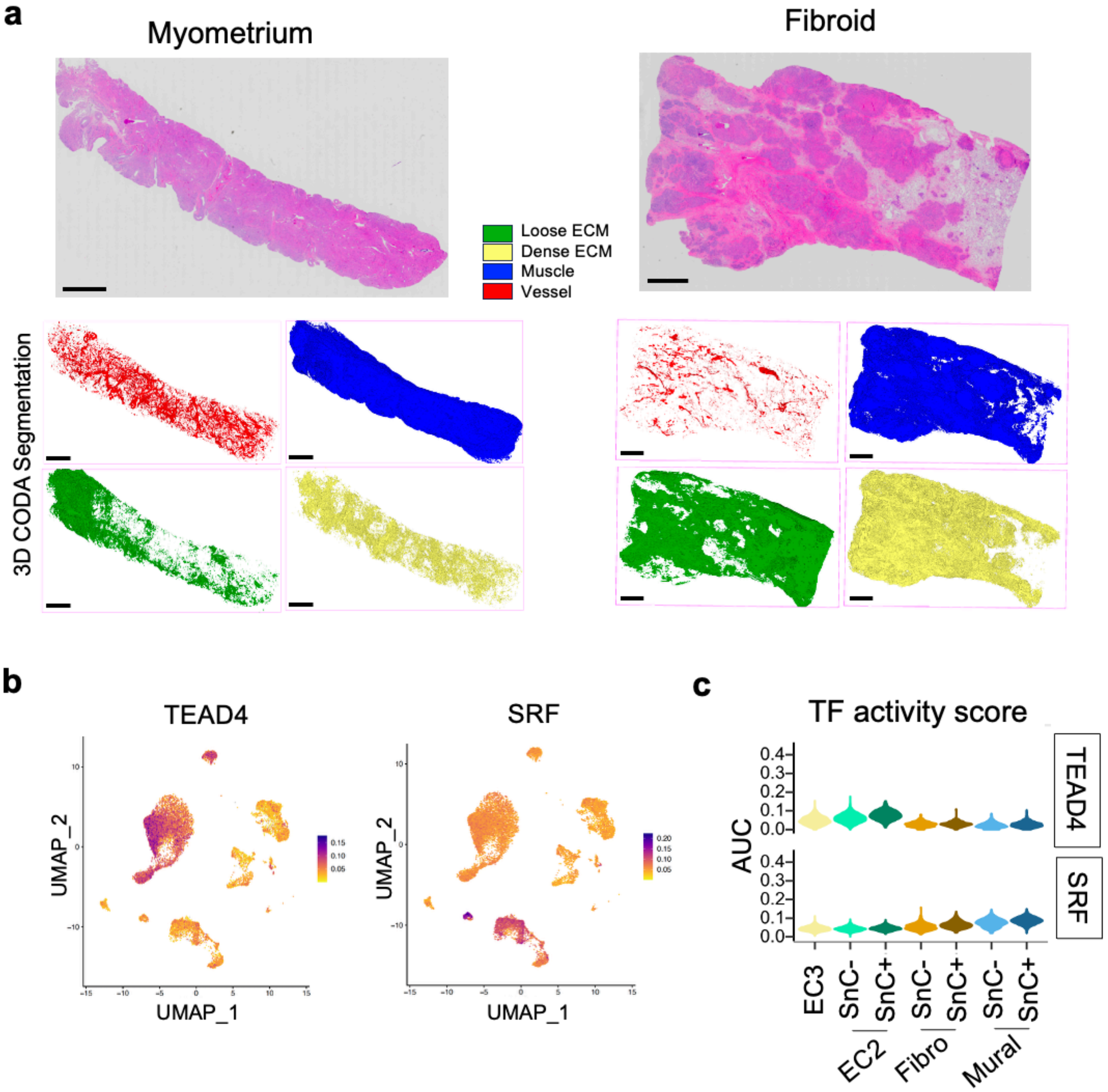


**Supplementary Fig. 17. ECM compartmentalization and mechanosensing regulon activity across fibroid cell states. (a)** Representative whole-section histological images of myometrium and fibroid tissue, together with corresponding CODA-derived tissue segmentation maps. Segmented compartments are shown as loose ECM (green), dense ECM (yellow), muscle (blue), and vascular structures (red). **(b)** UMAP feature plots showing single-cell *TEAD4* and *SRF* regulon activity across fibroid stromal and endothelial populations. Cells are colored according to SCENIC-derived regulon activity scores. **(c)** Violin plots comparing *TEAD4* and *SRF* regulon activity, quantified as area-under-the-curve (AUC) scores, across EC3 cells and SnC-negative and SnC-positive EC2, fibroblast, and mural populations.

**Supplementary Figure 18**

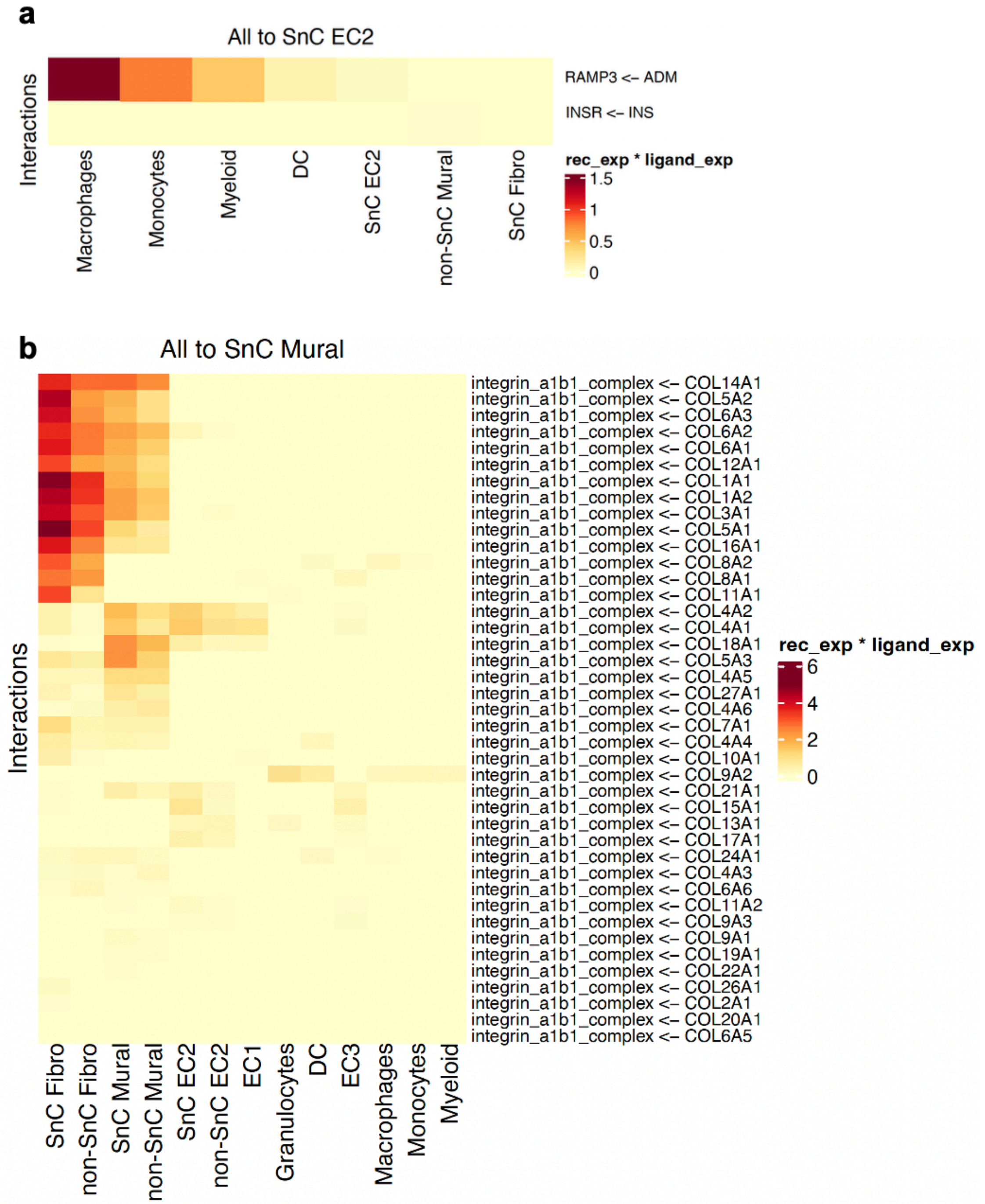


**Supplementary Fig. 18. Predicted incoming ligand–receptor signaling to SnC endothelial and mural populations. (a)** Heatmap showing predicted ligand–receptor interactions from the indicated sender populations to SnC EC2 endothelial cells. **(b)** Heatmap showing predicted incoming interactions to SnC mural cells, dominated by collagen ligands signaling through the integrin α1β1 receptor complex. Columns represent sending cell populations, and rows denote receptor–ligand pairs displayed as receptor ← ligand. Heatmap intensity represents the product of mean receptor expression in the receiving population and mean ligand expression in the sending population (rec_exp × ligand_exp). Color scales are specific to each panel.

**Supplementary Figure 19**

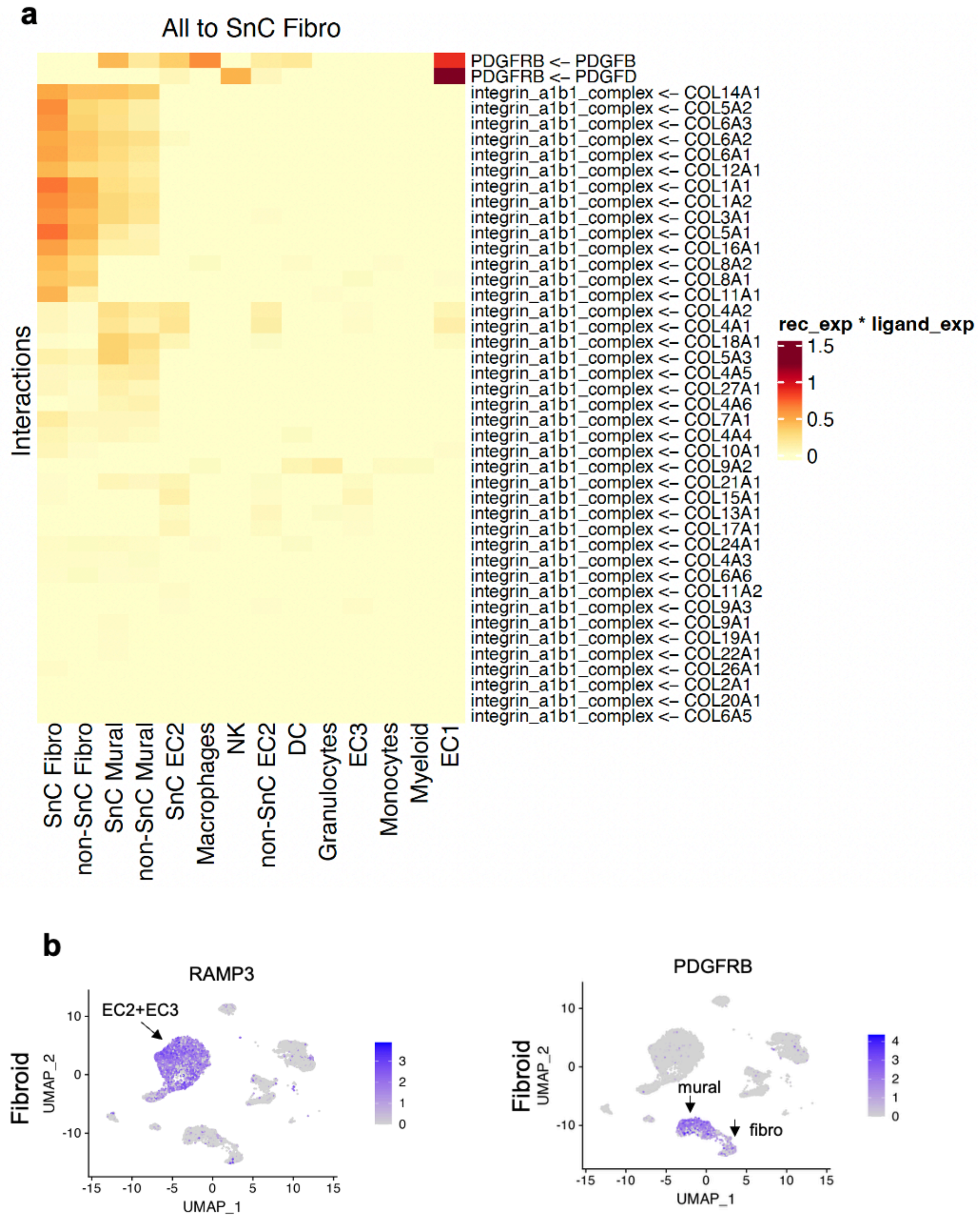


**Supplementary Fig. 19. Predicted incoming signaling to SnC fibroblasts and cellular localization of *RAMP3* and *PDGFRB* expression. (a)** Heatmap showing predicted ligand–receptor interactions from the indicated sender populations to SnC fibroblasts. The displayed interactions include *PDGFB–PDGFRB* and *PDGFD–PDGFRB* signaling and collagen ligands signaling through the *integrin α1β1* receptor complex. Columns represent sending cell populations, and rows denote receptor–ligand pairs displayed as receptor ← ligand. Heatmap intensity represents the product of mean receptor and ligand expression (rec_exp × ligand_exp). **(b** UMAP feature plots showing *RAMP3* and *PDGFRB* expression across fibroid cell populations. Arrows indicate enrichment of *RAMP3* expression in EC2 and EC3 endothelial populations and *PDGFRB* expression in mural and fibroblast populations.

**Supplementary Figure 20**

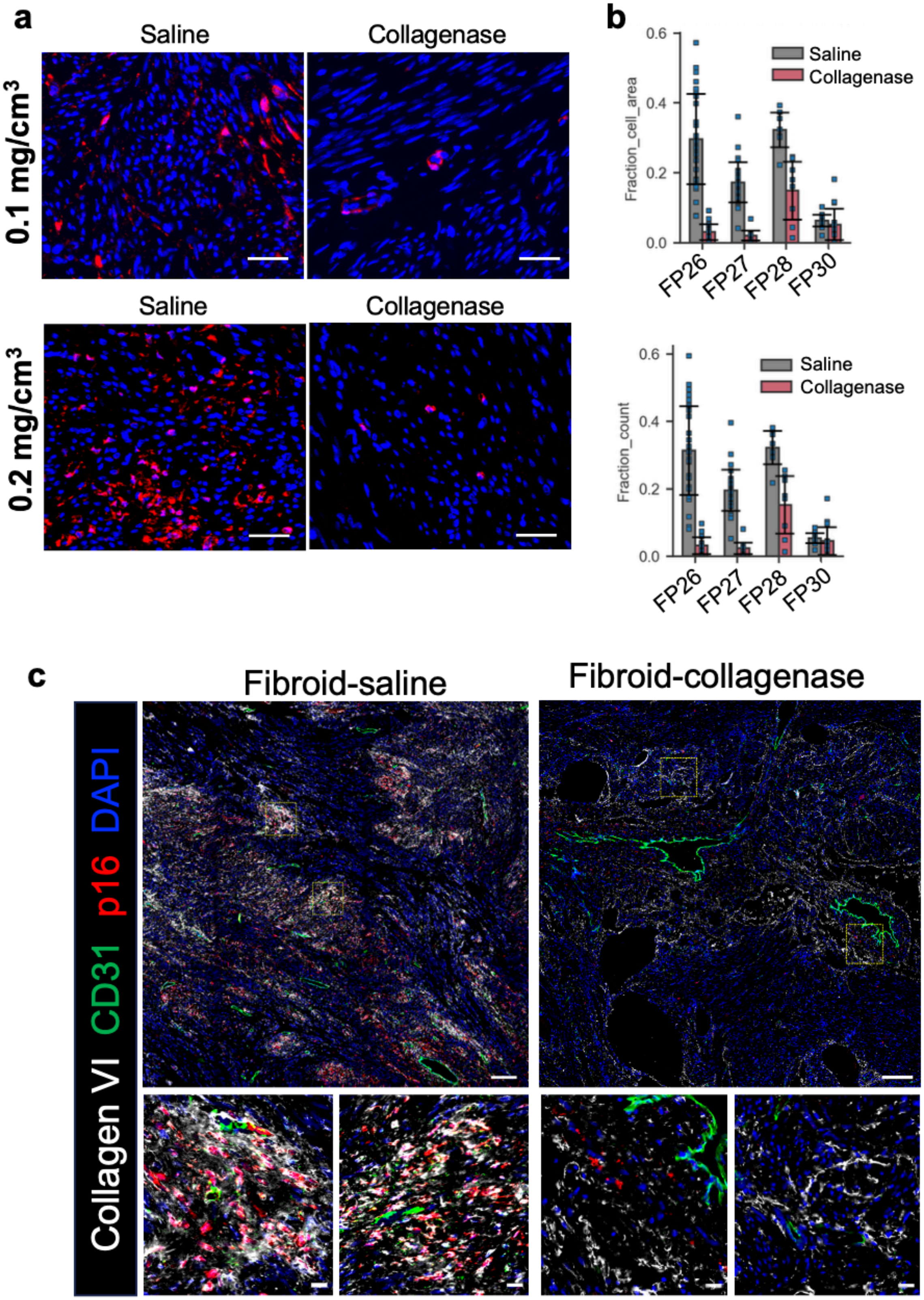

**Supplementary Fig. 20. Spatial and quantitative assessment of p16⁺ cells following collagenase treatment in uterine fibroids. (a)** Representative immunofluorescence images showing p16 (red) and DAPI (blue) in saline- and collagenase-treated fibroids from the 0.1 and 0.2 mg/cm³ dose groups. Scale bars, 50 µm**. (b)** Area- and count-based quantification of p16 positivity within spatial tissue subregions identified using the workflow shown in Fig. 6b. The upper graph shows the fraction of detected cell area assigned to $p16^+$ cells, and the lower graph shows the fraction of detected cells classified as $p16^+$. FP26 and FP27 received 0.1 mg/cm³ collagenase, whereas FP28 and FP30 received 0.2 mg/cm³ collagenase. Points represent individual tissue subregions; bars indicate the median and error bars indicate the interquartile range. **(c)** Whole-section multichannel immunofluorescence images of saline- and collagenase-treated fibroids from patient FP29, who received 0.2 mg/cm³ collagenase, showing DAPI (blue), collagen VI (white), CD31 (green) and p16 (red). Boxed regions in the whole-section images are shown at higher magnification below. Scale bars, 250 µm in the whole-section images and 20 µm in the enlarged regions.
